# A new chromosome-level genome assembly for western painted turtle *Chrysemys picta belli*, a model for extreme physiological adaptations

**DOI:** 10.1101/2025.04.01.646645

**Authors:** Junhao Chen, John R. Bermingham, Patrick Minx, Milinn Kremitzky, Zhenguo Lin, Daniel E. Warren

## Abstract

The western painted turtle, *Chrysemys picta bellii*, has the greatest tolerance to anoxia of any tetrapod studied to date. These turtles reside in the northern United States and southern Canada, and survive months of anoxia while submerged in ice-locked ponds and bogs. Reference genomes provide an important resource for elucidating the molecular bases for such unique physiological traits. An initial reference genome for this species was published in 2013, but the assembly is highly fragmented which poses several limitations for downstream analyses and biological interpretation. In this study, we created a new and improved assembly by combining PacBio HiFi, 10x Genomics Chromium, Hi-C sequence data and BioNano optical mapping derived from a single individual to generate a new haplotype-resolved chromosome-level assembly for *C. picta bellii*, called “SLU_Cpb5.0”. The genome size of the primary assembly is 2.372 Gb with a scaffold N50 of 133.6 Mb, which is a 6.5-fold improvement over the existing assembly. Genome annotation of SLU_Cpb5.0 revealed 12,242 novel genes compared to previous assemblies. Our PacBio Iso-Seq RNA sequencing data for twelve tissues unraveled over 100,000 novel transcript isoforms and 4,325 novel genes that were not annotated by standard NCBI pipeline. We also observed distinct patterns of tissue-specific isoform expression, creating a robust foundation for future characterization of the functions of these genes. The improved genome assembly and annotation will facilitate comparative genomics studies to better understand the genetic basis of *C.picta bellii*’s extreme physiological adaptations and other aspects of its biology.

## INTRODUCTION

Turtles are universally recognized because of their unique body plan, which results from the development of the shoulder girdle beneath fused ribs that form the dorsal carapace, formation of a ventral bony plastron, and replacement of teeth by a keratinous beak [1–4].

Additionally, they possess less conspicuous, but important specializations that make them fascinating and important research subjects, including long lifespans, environmentally determined sex determination, and the ability to tolerate environmental extremes, in particular low oxygen [5–7] and cold temperatures [8–11]. Notably, the western painted turtle (*Chrysemys picta bellii*, family Emydidae, **Additional Figure S1A**) has the greatest tolerance for anoxia (undetectable oxygen levels) of any tetrapod studied to date; as adults, they can survive anoxia for at least 30 hours at 20°C [12], and for more than 170 days at 3°C [5, 7].

*Chrysemys* (painted) turtles are polytypic, with a very broad North American distribution (**Additional Figure S1B**). Until recently *Chrysemys* turtles were described as having diversified into four subspecies, but analysis of mitochondrial DNA supports division of *Chrysemys* into two clades, with Southern painted turtles sufficiently diverged to be considered a distinct species (*C. dorsalis*), and with *C. picta* consisting of three subspecies, Eastern (*Chrysemys picta picta*), Midland (*C. picta marginata*), and Western (*C. picta bellii*) [13, 14]. The *C. picta bellii* range exceeds those of the other *Chrysemys* taxa combined; currently it extends from northern Texas northward into southern Manitoba, and westward into Oregon, Washington, and southern British Columbia, with relict populations in New Mexico [15, 16]. *C. p. bellii’*s unique tolerance of anoxia, a critical adaptation to survive long winters, likely facilitated the species’ range expansion.

The extreme hypoxia tolerance of painted turtles was first documented decades ago [5, 17]. Work since then has shown that survival relies on the avoidance of cellular ATP depletion during the anoxic bout [18–20], and preventing damage from reactive oxygen species (ROS) during reoxygenation (i.e., reperfusion injury; [21, 22]), the two principal causes of cellular injury in anoxia-sensitive animals, including humans [23, 24]. The first is achieved through a temperature-independent metabolic suppression of more than 90% [25], enabling ATP requirements to be met entirely through glycolysis, which is supported by the species’ unusually large tissue glycogen stores [26, 27], and its highly carbonated skeletal mass, which prevents lethal decreases in body fluid pH caused by extreme lactic acidosis [6, 7]. The second is achieved primarily through the avoidance of mitochondrial ROS production [21, 22]. Many of the underlying mechanisms remain a mystery, and an improved western painted turtle genomic resource should facilitate their further characterization, potentially leading to the identification of new targets for therapeutic intervention in the treatment of ischemia-related conditions in humans associated with stroke, cardiac arrest, and organ transplantation.

The genome of the western painted turtle was the first turtle assembly to be published [28], and as such, has been cited more than any other turtle genome study. Despite recent improvements [29, 30], however, this original assembly remains highly fragmented because it was generated using a combination of 454 technology, and Illumina short reads, compromising its utility for trait mapping and gene discovery. Furthermore, DNA that was used for the original assembly was obtained from two turtles in the state of Washington, at the northwestern margin of the species’ geographic range, one of which was used for sequencing, and the second for assembling a bacterial artificial chromosome (BAC) library. Subsequent BioNano optical maps were derived from a third turtle of uncertain origins [30]. These turtles may possess single nucleotide polymorphism (SNP) or structural differences with each other, and with turtles from the center of the range. For these reasons, we generated a new reference genome for this species that is consistent with standards that have been established by the Vertebrate Genome Project (VGP; [31]). We used tissue from the same individual (Biosample SAMN13293682), collected from McLeod County, MN near the center of the species’ range (**Additional Figure S1B**), to generate datasets using 10XGenomics, PacBio HiFi, and Hi-C sequencing, as well as BioNano optical maps; all four datasets were used to generate new *Chrysemys picta bellii* genome assemblies (haplotype1, haplotype2, merged). To annotate our assembly(ies), we generated PacBio IsoSeq transcriptomes from twelve tissues, to identify as much transcript diversity as practical [32]. Additionally, we utilized previously generated Illumina short-read RNAseq datasets from brain and heart of normoxic and anoxic adult and hatchling turtles acclimated to different temperatures [33, 34]. Collectively, these transcriptomes allow a more complete annotation of our new *C. picta bellii* reference genome than those currently available for other turtle genomes. The chromosomal level assembly and annotations for *C. picta bellii* generated in this study will facilitate future studies on the unique physiology and evolutionary biology of the western painted turtle, as well as their phylogeny.

## RESULTS

### *De novo* assembly of SLU_Cpb5.0, a new *Chrysemys picta bellii* genome

Initially we generated a new *C. picta bellii* genome assembly from 10x Genomics linked reads [35] – 572.7 million pairs of Illumina reads (2 × 150 bp), with a total length of 171.8 Gb. This 10x Genomics assembly (ASM1138683v1; referred as SLU_Cpb1.0 **Additional Table1**; **Additional Figure S2**) consisted of 132,827 scaffolds with a Scaffold N50 of 15.9 Mb. To construct an improved, chromosome-level *C. picta bellii* assembly, we generated additional PacBio HiFi long reads, Hi-C conformation capture sequence data, and Bionano optical genome maps from the same individual turtle (See Materials and Methods, **Additional Figure S2**). We obtained a total of 3.48 million HiFi reads from three PacBio SMRT cells (**Additional Table S2, Additional Figure S3**). The mean HiFi read lengths range from 13,876 to 14,024 bp, with median read Phred quality scores between 34 and 35 (**Additional Table S2**). The total length of HiFi reads was 48.5 Gb, providing an average coverage of ∼ 20x based on the previously assembled genome size of 2.4 Gb (Shaffer et al. 2013). Our Hi-C sequencing produced 496.7 million pairs of Illumina reads (2 × 150 bp) (**Additional Table S2**).

To achieve the best assembly quality, we tried different *de novo* genome assembly tools that provide haplotype-resolved assemblies, including Hifiasm [36], NextDenovo [37], and Wtdbg2 [38]. We noted that some assemblies with better contiguity statistics could possess incorrectly merged contigs (Salzberg and Yorke 2005). We found that Hifiasm, with integration of Hi-C reads for haplotype phasing, yielded the best metrics in most categories for primary, haplotype1, and haplotype2 preliminary assemblies (assembly version SLU_Cpb2.0, Contig N50 size: primary assembly = 28.07 Mb, haplotype 1 = 3.54 Mb, and haplotype 2 = 3.57 Mb; **Additional Table S3, Additional Figure S2**). Thus, the assemblies that combined PacBio HiFi + Hi-C datasets by Hifiasm were used for subsequent scaffoldings.

The first round of scaffolding for the preliminary assemblies was performed with the 10x Genomics linked reads using Tigmint [39]. The scaffold N50 size for the primary assembly was improved to 42.12 Mb, and 38.31 Mb and 43.67 Mb for haplotype 1 and 2, respectively (assembly version SLU_Cpb3.0, **Additional Table S3**). Hi-C sequencing reads were used for the second round of scaffolding by 3D-DNA [40], which linked scaffolds into 25 pseudochromosomes (assembled sequences that correspond to actual chromosomes; assembly version SLU_Cpb4.0, **Additional Figure S4**). Hi-C scaffolding significantly improved the contiguity of all assemblies, as scaffold N50 sizes for the primary assembly, haplotype 1 and haplotype 2 were increased to 133.42 Mb, 132.97 Mb, and 132.08 Mb, respectively. All these N50 values are a significant improvement to the 20.68 Mb of the previous assembly (NCBI RefSeq assembly GCF_000241765.5 “Chrysemys_picta_BioNano-3.0.4”, abbreviated as CPI3.0.4 hereafter) (**Table 1**). The primary assembly was aligned with BioNano optical maps, and major discrepancies were corrected manually (**Additional Figures S2 and S5**).

**Table 1:**
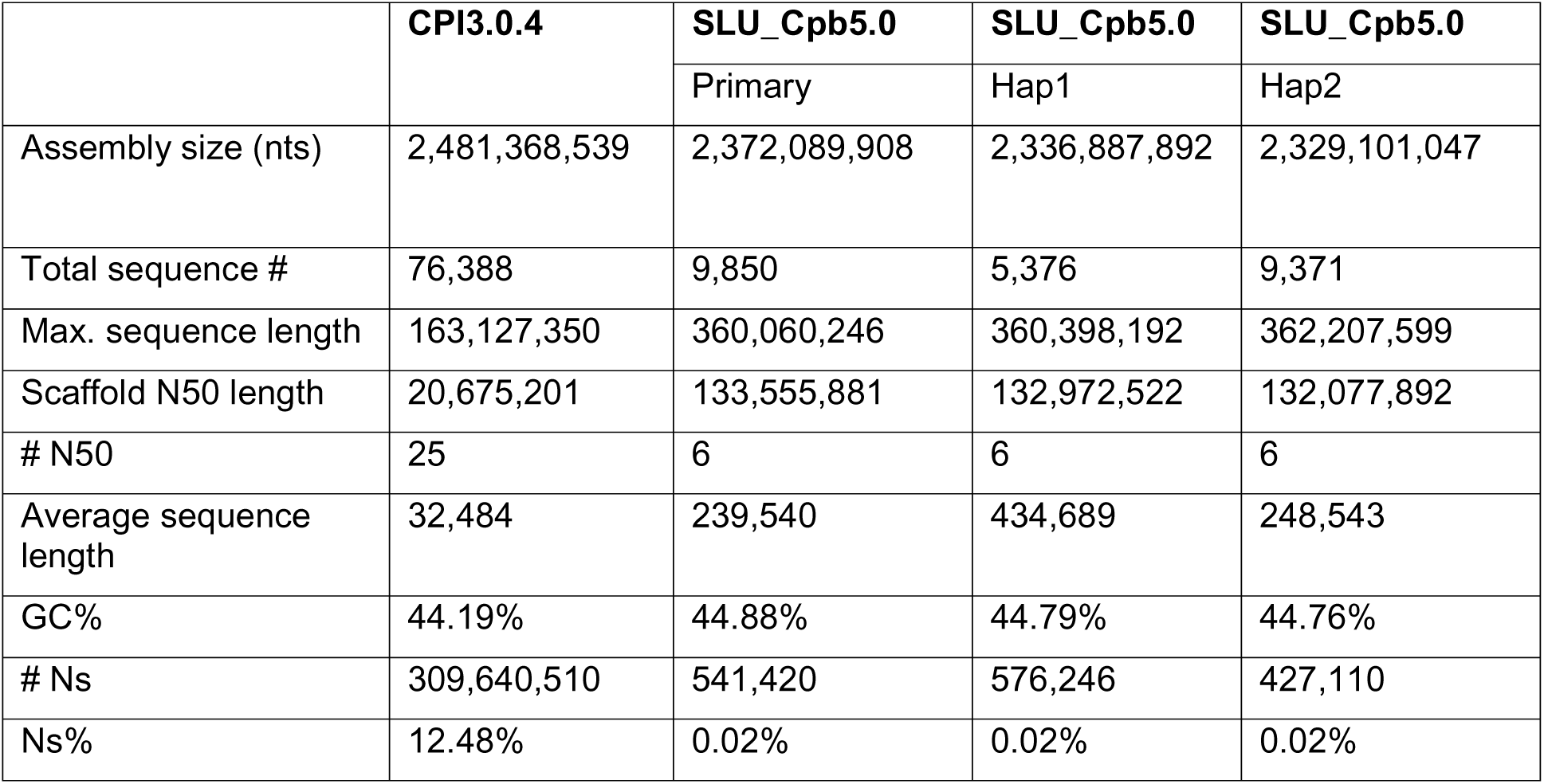
Comparison of summary statistics for previous and new *Chrysemys picta bellii* primary and haplotype assemblies. The table compares assembly statistics for the older CPI3.0.4 assembly [28, 29] and the SLU_Cpb5.0 assembly. PacBio HiFi and Hi-C sequencing generate both a primary assembly and haplotype-resolved assemblies. #N50 is the number of scaffolds, starting with the largest, that are required to cover 50% of the genome. #Ns refer to unknown bases in the assemblies.

Our final primary assembly version “SLU_Cpb5.0”, has a total size of 2.372 Gb, which closely matches the estimated genome size of 2.314 Gb based on k-mer analysis using10x Genomics Illumina reads (**Additional Figure S6A**), but is larger than 2.102 Gb inferred using PacBio HiFi reads (**Additional Figure S6B**). The difference in estimated genome size based on k-mer analysis between 10x Genomics Illumina reads and PacBio HiFi long reads is likely because our 10x Genomics data have a much higher sequencing depth (72.5× vs 20.5×), and a higher sequencing depth generally provides a more accurate estimation of genome size in k-mer analyses [41].

The primary and haplotype assemblies consist of 25 nuclear pseudochromosome sequences (placed scaffolds), corresponding to the species’ 25 chromosomes. As chromosomes are conventionally numbered according to size, the pseudochromosomes from our assembly are numbered from largest to smallest accordingly **(Figure 1A)**. The total number of unplaced scaffold sequences has significantly improved, with SLU_Cpb5.0 having only 9,850, compared to 76,388 in the previous assembly CPI3.0.4. In addition, the average scaffold sequence length was increased by 7.4x, the percentage of Ns was reduced from 12.48% to 0.02%, which partially explains the smaller estimated size of the current assembly (2.372 Gb vs. 2.481 Gb) (**Table 1)**.

**Figure 1:**
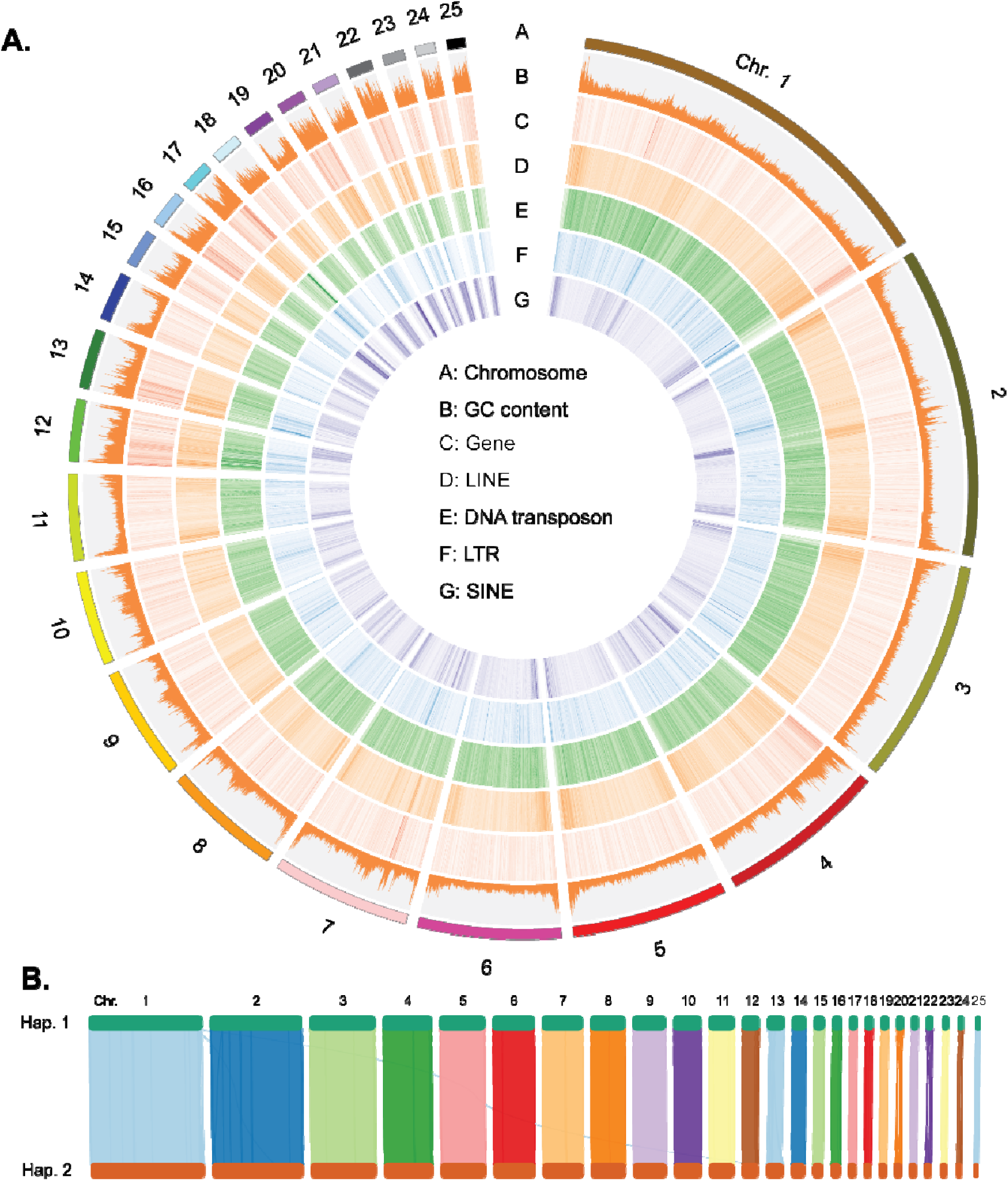
Structure of primary assembly of the *Chrysemys picta bellii* genome SLU_Cpb5.0. **A.** Circos plot of the 25 pseudochromosomes of the *C.picta bellii* genome. Different genomic features are depicted in seven concentric circles. The outer circle, labeled A, depicts the 25 assembled pseudochromosomes, which correspond to cytological chromosomes. Circles B-C: GC content and gene density, respectively. GC content increases toward telomeres and perhaps near some centromeres, and is increased in the smaller microchromosomes. Circles D-G represent densities of repetitive elements: LINEs, DNA transposons, LTRsand SINEs. Diagrams of the assembled genome were generated using Circos [128], and diagrams of collinearity between two haplotype assemblies, and between closely related turtle genomes, were created using NGenomeSyn [129]. **B.** Collinearity between the 25 chromosomal pairs of haplotypes 1 and 2. Each line between the haplotypes represents a colinear block of 100 kb. The two haplotypes are largely colinear; subtle differences may reflect heterozygous indels or could be assembly errors.

To assess the structural and sequence-level consistency between the two haplotypes, we conducted collinearity analysis between the haplotype 1 and haplotype 2 assemblies. As shown in **Figure 1B**, the two haplotypes align well with one another and are consistent in terms of large-scale genomic structure. The mean heterozygosity rate between the two haplotypes is 0.48%, which is consistent with 0.548% based on 10x Genomics Illumina reads using k-mer analysis by GenomeScope 2.0 [41], but is lower than 0.712% estimated using PacBio HiFi raw reads (See Material and Methods, (**Additional Table S3, and Figures S5**). The discrepancy in estimated heterozygosity between the -two types of sequencing data is likely attributed to their different sequencing depths, or to poorer alignment of short reads at polymorphic sites. Nevertheless, all these numbers suggest high levels of genetic diversity in Minnesota western painted turtles when compared to individuals of different turtle species, as well as to the *C. picta bellii* individual from Washington state [42].

### The new assembly SLU_Cpb5.0 is a significant improvement over the previous assembly CPI3.0.4 for the *C. picta bellii* genome

The completeness of the primary and haplotype assemblies was evaluated with the BUSCO v5 using the sauropsida_odb10 dataset [43]. As shown in **Figure 2A**, the primary assembly achieved a BUSCO completeness score of 99.1%, indicating near-complete recovery of single-copy orthologs.

**Figure 2.**
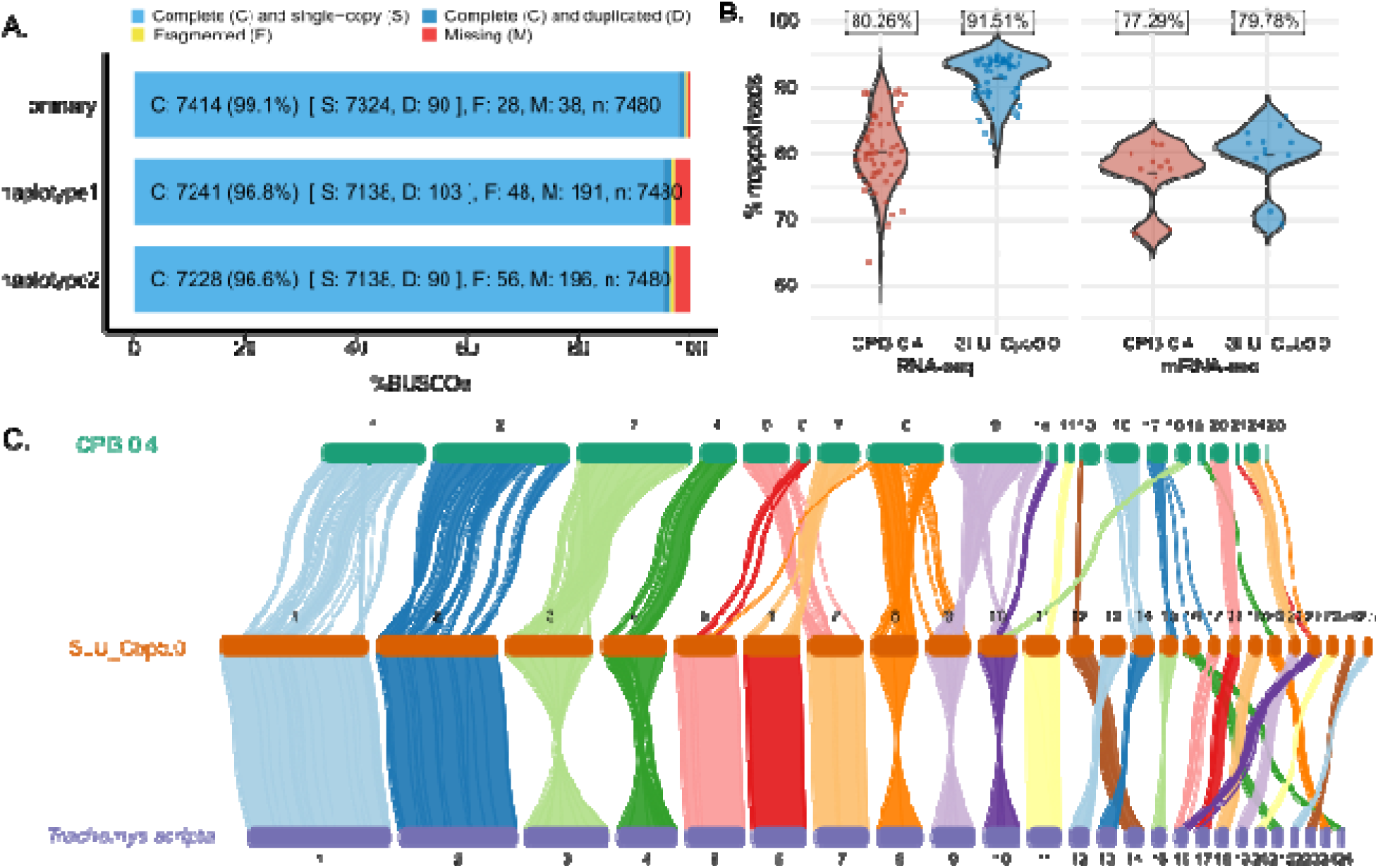
Evaluation of the SLU_Cpb5.0 assembly(ies) of the *C. picta bellii* genome. A. BUSCO plot for the primary (merged) genome, haplotype 1, and haplotype 2. The results are categorized into four groups: Complete (C), which includes single-copy (S) and duplicated (D) orthologs; Fragmented (F), representing partially recovered orthologs; and Missing (M), indicating orthologs absent in the dataset. The number of genes and percentages are shown for each category. These results are based on comparison with the sauropsida_odb10 lineage dataset. B. Improved alignment rates of RNAseq reads to new *C.picta bellii* genome assembly. Violin plots display the alignment rates for RNAseq datasets. Each dot is a dataset, and it is plotted according to the percentage of its reads that align to each genome assembly. For each violin plot, the median alignment percentage is depicted by a horizontal bar, and is listed at the top. The percentage of reads aligned for mRNAs (left) and microRNAs (right) is better for the SLU_Cpb5.0 assembly than for the 3.0.4 assembly. Increasing assembly quality has a greater impact on mapping efficiency of mRNAs than it does for the smaller microRNAs. C. Pseudochromosome homologies between *Chrysemys picta bellii* and *Trachemys scripta* assemblies. Alignment of the *C. picta bellii* assembly SLU_Cpb5.0 with assembly CPI3.0.4 and with the red-eared slider genome assembly CAS_Tse_1.0 [45], showing substantial contiguity between the two chromosome-level assemblies. The figure was generated using NGenomeSyn [129], with a 10Kb cutoff for homologies.

The mapping rate of RNA-seq reads can be used as a useful metric to evaluate the quality of a genome assembly, particularly in terms of completeness. To determine the extent to which our new assembly improves RNA-seq mapping rate, we downloaded 59 RNA-seq datasets of *C. picta bellii* from the NCBI SRA database (**Additional Table S5**). These RNA-seq datasets were mapped to both SLU_Cpb5.0 and the CPI3.0.4 assemblies using HISAT2 [44]. Our results show that the RNA-seq mapping rate using the SLU_Cpb5.0 assembly is significantly higher than that for the 3.0.4 assembly (median 91.32% vs. 80.25%, **Figure 2B**), demonstrating that our assembly captures more coding regions in the genome and has a lower number of assembly errors and misassembled regions. Interestingly, the improvement in mapping rates for microRNA sequencing (miRNA-seq) was modest compared with those for regular RNA-seq data (**Figure 2B)**, suggesting that the smaller miRNA genes are less sensitive to fragmented genome assemblies than are the longer mRNAs.

Aligning assembled sequences to those of a closely related species also reveals insights into the accuracy, completeness, and structural integrity of the assembly. We aligned our assembly to the reference genome of a closely related Deirochelid species *Trachemys scripta* (NCBI RefSeq assembly GCF_013100865.1 “CAS_Tse_1.0”), which also contains 25 chromosomes [45]. As shown in **Figure 2C**, the lengths of 25 chromosomes are highly similar between SLU_Cpb5.0 and *T. scripta* “CAS_Tse_1.0”. In contrast, the assembled chromosomes in the CPI3.0.4 assembly are significantly shorter than the other two assemblies, reflecting the many unmapped scaffolds in the CPI3.0.4 assembly. In addition, higher degrees of collinearity are observed in orthologous chromosomes between SLU_Cpb5.0 and *T. scripta*, than between SLU_Cpb5.0 and the CPI3.0.4 assembly, particularly from chromosome 4 to chromosome 10 (**Figure 2C**). These results demonstrate that the CPI3.0.4 assembly is not only highly fragmented, but also misassembled across many chromosomes.

Bacterial Artificial Chromosomes (BACs) can serve as valuable tools to evaluate genome assembly quality. Many *C. picta bellii* BACs have been mapped to chromosomes by fluorescence in situ hybridization (FISH; [29, 30]). We localized those BACs that have been assigned NCBI accession numbers to our SLU_Cpb5.0 assembly, to correlate the chromosomes that have been labelled by FISH to the pseudochromosomes that have been generated by our assembly. Our BAC localizations demonstrate the improved contiguity of our assembly (**Additional Figure S7)**. Cytogenetic chromosome and sequence-based pseudochromosome numbers are listed in **Additional Table S6.** As we numbered the pseudochromosomes from our assembly according to size, the sizes of our pseudochromosomes are largely consistent with chromosome numbers that were assigned based on cytology [29, 30]. However, some differences are apparent: Pseudochromosomes 5, 6, and 7 contain BAC sequences that were hybridized to cytologically numbered chromosomes 7, 5, and 6, respectively. Similarly, numerous smaller pseudochromosomes contain BAC sequences that hybridize to different cytologically numbered chromosomes. We suggest that the very similar sizes of compact metaphase chromosomes on the slides led to their misnumbering. Because SLU_Cpb5.0 assembly is not a complete telomere-to-telomere assembly [46], however, unassembled centromere and telomere repetitive sequences could affect pseudochromosome numbering that is based on sequence length.

Two BACs, CHY3-12H12 and CHY3-34H12, appear to have been mis-localized by FISH. Badenhorst (2015) and Lee (2020) list CHY3-12H12 as residing on chromosome 19 of *C. picta bellii* and *Trachemys scripta* [29, 30], and additionally, Bista (2024) lists this BAC as residing on chromosome 20 of *Saurotypus triporcatus,* the Mexican musk turtle [47]. However, its sequence (NCBI accession AC239503.2) resides on pseudochromosome 3 in the SLU_Cpb5.0 assembly. BLAST searches of Whole Genome Sequence datasets for *Trachemys scripta*, *Saurotypus triporcatus*, and *Mauremys reevesi* genomes confirmed that AC239503.2 sequences reside on chromosome/linkage group 3. The second BAC, CHY3-34H12 (accession AC238643.2) was listed in Badenhorst (2015) as residing on *C.picta bellii* chromosome 1p, whereas its sequence covers 195.9-196.1M on pseudochromomsome 3 in the SLU_Cpb5.0 assembly. BLAST searches of *Trachemys scripta, Saurotypus triporcatus*, and *Mauremys reevesi* genomes also localize AC238643.2 sequences to chromosome3/linkage group 3. We believe that our placement of these BACs is correct for two reasons. First, it is extremely unlikely that independent assemblies of four different turtle genomes all misassembled this sequence. Second, our *C. picta bellii* BioNano optical map of this region shows no evidence of mis-assembly (data not shown). These observations demonstrate the utility of correlating whole genome sequences and FISH results, and will improve the reliability of future turtle cytological studies.

### Comparative studies of mitochondrial genomes reveal rapidly evolving regions

Mitochondria play a central role in energy metabolism and metabolic signaling (reviewed in [48], including two traits of interest in painted turtles: anoxia tolerance and slow aging. Furthermore, mitochondrial polymorphisms have been used to address questions about *Chrysemys* taxonomy and population stratification [49]. We assembled the mitochondrial genome of *C. picta bellii* using our HiFi reads using MitoHiFi [50], which created a circular sequence with a length of 16,931bp. Annotation of the mitochondria genome using MITOS2 [51] predicts 13 protein-coding genes, 22tRNA genes and 4 rRNA genes.

We examined differences in *Chrysemys* mitochondrial genomes by aligning our mitochondrial genome with the four other publicly available *Chrysemys* mitochondrial genomes: *C. dorsalis* (introduced into Korea, original provenance unknown [52]), *C. picta picta* (provenance unknown), *C. picta bellii* (Grant Co. WA; [53] – sequences obtained from [28]), and *C. picta bellii* (also introduced into Korea, original provenance unknown; [54]). Pairwise percent identities between the genomes are listed in **Additional Table S7;** they range from 98.46% (*C.picta bellii* (WA) *vs C. dorsalis*) to 99.92% (*C. picta bellii* (Korea) to the assembly that is presented here, *C. picta bellii* (MN)). We observed variation in mitochondrial tRNA, rRNA, and protein coding genes, but most variation occurred in noncoding regions of the mitochondrial genome. A summary of mitochondrial variation between the three *C. picta bellii* mitochondrial genomes is presented in **Additional Table S8.** These variants may contribute to physiological differences between *C. dorsalis* and *C.picta bellii* [55].

Additionally, mitochondrial differences can inform estimates of population divergence. The differences that we observe between *C. picta* and *C. dorsalis* mitochondrial sequences, 1.21-1.54% (**Additional Table S7**), is comparable to the 1.27% divergence between the human rCRS mitochondrial genome (accession NC_012920.1) and a Neanderthal mitochondrial genome NC_011137.1 [56]. Estimates for the time to most recent common ancestor (TMRCA) for humans and Neanderthals range from 400 thousand years (kya) to 800 kya [57]; given that the per-year mitochondrial mutation rate is approximately 2.69× faster in humans (0.01665 mutations/bp/10^6^ years; [58]) than in turtles of the superfamily Testudinoidea (which includes *Chrysemys*; 0.00618 mutations/bp/10^6^ years; [59], we calculate that the *C. picta*-*C. dorsalis* split occurred 1-2 Mya, supporting the proposed designation of *C. dorsalis* as a separate species. Similarly, our painted turtle mitochondrial genome also permits an estimate of divergence times between *Chrysemys* populations. Given the *C. dorsalis*-*C.picta* divergence time estimate, the 0.36% difference between Minnesota and Washington *C. picta bellii* mitochondrial genomes suggests that these *C. picta belli* populations diverged 300,000-600,000 years ago.

The greatest differences between the *Chrysemys* mitochondrial genomes were found in two regions: A GC-rich region between the 3’ ends of the oppositely transcribed genes mt-ND5 and mt-ND6 varied between all five *Chrysemys* mitochondrial genomes, and may be useful for future phylogeographic studies. The second is the well-studied Control Region, which contains the D-loop, and mitochondrial origins of replication and transcription. In turtles, the mitochondrial Control Region consists of Left, Central, and Right Domains [60]. The Left Domain contains termination-associated sequences, and for some turtles, a variable number of tandem repeats (VNTR’s). We found that the *C.picta picta* and the two previously published *C. picta bellii* mitochondrial genomes possess a Left Domain VNTR that consists of two 59 bp VNTR1A repeats flanking a 33 bp C-rich VNTR1B repeat (an “A-B-A” configuration) whereas the *C. picta bellii* mitochondrial genome reported here has additional 59 bp and 33 bp repeats (“A-B-A-B-A”; **Additional Figure 8A**), similar to the *Trachemys scripta* mitochondrial genome. The *C. dorsalis* Left Domain VNTR possesses five 33 bp repeats interspersed between six 59 bp repeats (“A-B-A-B-A-B-A-B-A-B-A”). A potential explanation for the smaller number of *C. picta* VNTR1 repeats, but not for the expanded numbers in the *C. dorsalis* VNTR1 repeat, is the difference in sequencing methodology. Short-read sequencing of mitochondrial genomes tends to artificially delete repeats [61].

We searched for sequences similar to VNTR1A and VNTR1B because Bernacki et al. [60] reported Left Domain VNTRs only in Trionychidae and *Trachemy*s. A BLASTn search for sequences similar to the *C. picta bellii* VNTR1A, produced the following observations: 1) VNTR1A is present in mitochondrial DNA from multiple Emydid genera, including *Emys, Pseudemys, Graptemys, Malaclemys* and *Trachemys* (**Additional Figure 8B**). 2) Different numbers of VNTR1A repeats are observed in different *Emys orbicularis* sequences, demonstrating that VNTR1A expansion occurs within this species as well. 3) All six *C. dorsalis* VNTR1A repeats have the same pair of polymorphisms, suggesting that they expanded from a single diverged VNTR1A repeat. 4) The observation of VNTR1A repeats in *Emys* and *Malaclemys* with diverged sequences suggests that they have begun to accumulate mutations, and that the expansions in *Chrysemys* may be evolutionarily recent. Similarly, *C. picta bellii* VNTR1B homology is detectable by BLASTn only in mitochondrial sequences from *Chrysemys* turtles, but extension of the VNTR1B sequence flanked by additional ≥13nt revealed homologies to additional turtles belonging to the Emydid subfamily Deirochelyinae. Examination of alignments with European pond turtle, Emys orbicularis, subfamily Emydinae, reveals that the *Emys* VNTR1B is highly diverged, suggesting that it is more ancient than the VNTR1A domain, or that it is diverging more rapidly. We conclude that VNTR1 repeats are likely present in all Emydid turtles, and are evolving rapidly.

### Annotation of the *C. picta bellii* SLU_Cpb5.0 genome assembly

Annotation of the primary assembly was completed by the NCBI eukaryotic genome annotation pipeline (see Materials and Methods) [62]. The final annotation yielded 35,699 genes, including 25,216 protein-coding genes and 1,155 pseudogenes. In terms of non-coding RNA, the annotation identified 219 small nuclear RNA (snRNA) genes, 283 small nucleolar RNAs (snoRNAs) genes, 1,269 ribosomal RNA (rRNA) genes, 468 transfer RNA (tRNA) genes, and 6965 long non-coding RNA (lncRNA) genes. The genome coverage analysis showed that 5.27% of the assembly is covered by exons, while introns account for 43.51%. Among the protein-coding genes, 20,705 (82.1%) were assigned to at least one Gene Ontology (GO) term, suggesting potential function(s). At the transcript level, a total of 114,986 transcripts were annotated, with an average of 3.22 transcripts per gene. Among them, 85,016 are mRNA transcripts, spanning lengths from 156 to 114,607 bp.

Transposable elements and repetitive sequences were inferred from the primary assembly using Earl Grey with default parameters [63]. In terms of size, the top four largest groups of repetitive sequences are LINE, DNA transposon, LTR and SINE, which account for 19.07%, 9.58%, 5.2%, 2.01% of the turtle genome, respectively **(Figure 3A).** The proportions of SINE and LTR sequences we observed are similar to those observed in the original assembly of western painted turtle [28]. However, a higher proportion of LINE elements were identified in this assembly, probably due to improved genome continuity.

**Figure 3:**
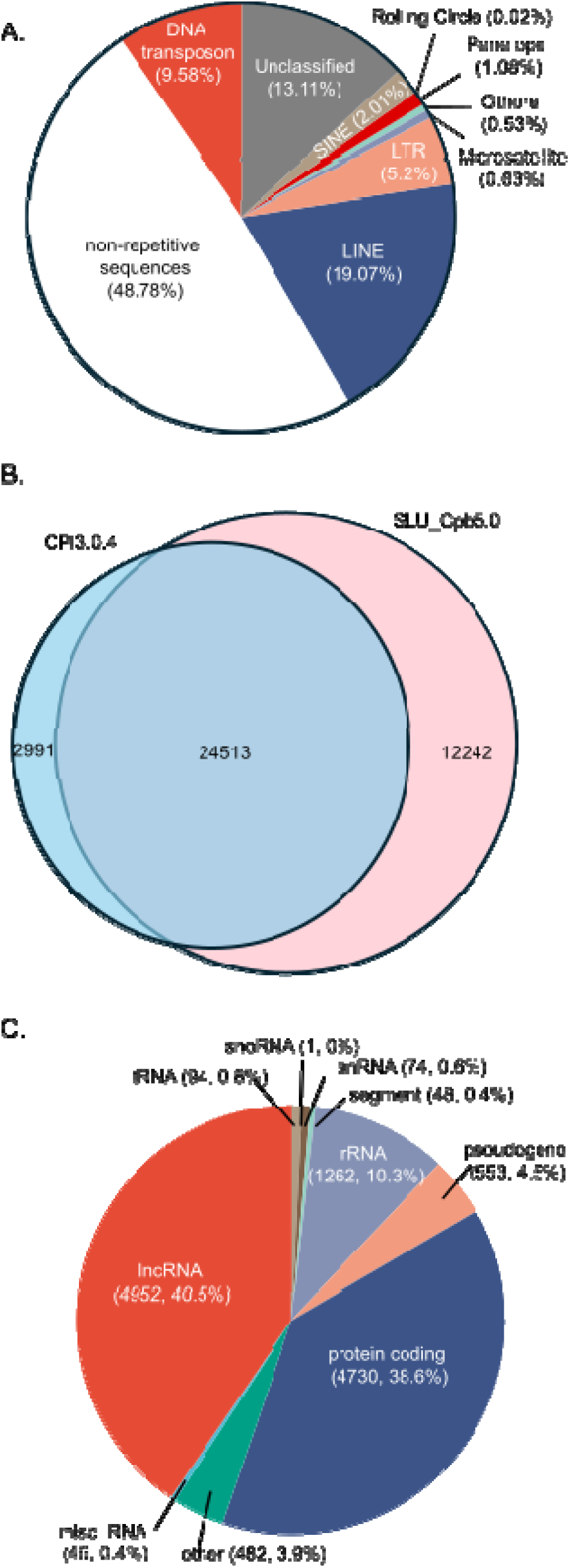
Annotations of the *C. picta bellii* genome. **A.** A pie chart showing the distribution of different types of repetitive elements in the *C. picta bellii* genome. **B.** Venn diagram showing the differences in gene annotation between two different assemblies of the *C. picta bellii* genome (CPI3.0.4 vs. SLU_Cpb_2.0). **C.** A pie chart showing the distributions of different types of genes that were annotated only in the new assembly SLU_Cpb_2.0.

When we compared the annotations between the two assemblies, SLU_Cpb5.0 annotations include 12,242 genes that are absent in CPI3.0.4 **(Figure 3B).** On the other hand, 2,991 genes annotated in CPI3.0.4 are no longer present in the SLU_Cpb5.0; this observation likely results from an increase in stringency of annotation parameters in the last decade, leading to the exclusion of some previously annotated candidate genes from our annotation. Among the newly predicted genes, 4,952, (40.5%) are annotated as long noncoding RNAs), and 4,730 (38.6%) as protein-coding genes (**Figure 3C**). The third largest group of newly annotated genes are rRNA genes (1,262). This is notable because eukaryotic genomes typically contain numerous rRNA loci with high sequence identities, which poses significant challenges for assembly with short-read sequencing technologies. By utilizing PacBio long reads, we were able to achieve a substantial improvement in the assembly of these repetitive rRNA loci.

### Identification of tissue-specific transcript isoforms in western painted turtle using PacBio Iso-Seq data

We generated PacBio Iso-Seq long-read transcript datasets using 12 tissues from adult and hatchling western painted turtles (**Additional Table S9**). These tissues were chosen either because they are expected to express a large fraction of the genome (testis), are important for anoxia/cold tolerance (hatchling shell, adult and hatchling kidney, adult liver, adult skeletal muscle, adult hindbrain, and hatchling brain), or because they are underrepresented in other painted turtle transcript datasets (adult lung, adult spinal cord, adult sciatic nerve, and adult duodenum) and would, therefore, expand the completeness of gene identification in our assembly. Alternative splicing is prevalent in vertebrates, and the expression of many transcript isoforms is very often restricted to specific tissues. [64]. These Iso-Seq datasets also allow us to determine the extent to which alternative splicing occurs in the western painted turtle, and to identify tissue-specific transcript isoforms.

The transcript models in each examined tissue were constructed by mapping Iso-Seq reads to assembly SLU_Cpb5.0. The splice junctions of transcript models were further validated by using public short read RNA-seq data (see Methods and Materials). Each transcript construct was compared to NCBI annotations to identify novel transcript isoforms and genes. Although Iso-Seq data are highly effective in capturing full-length transcripts, they have limitations in reliably defining the 5’ and 3’ ends of transcripts due to several technical factors [65]. To exclude potential false novel transcript isoforms, we applied stringent criteria to define novel transcript isoforms by excluding those with altered 5’ and 3’ ends but splice junctions are fully matched with NCBI annotated transcripts, as well as those with incomplete splice junction matches (potential partial transcripts).

In general, the number of transcript isoforms and expressed genes differ significantly from tissue to tissue (**Figure 4A**, **Additional Table S10**). Our results show that hatchling carapace has the most expressed genes detected, following by testis. Although we expected that testis expressed many genes, our observation of more expressed genes in hatchling carapace was unexpected. It may be because carapace is a mix of many different tissues, such as blood, muscle, bone, connective tissues, epidermis (scutes). The hatchling kidney and adult sciatic nerve have the least number of expressed genes and transcript isoforms. A likely explanation is that the input total RNA from these tissues was much less for these than the other tissues, resulting in a much lower Iso-Seq output (**Additional Table S9**). For hatchling kidney, another possibility that hatchling painted turtles do not utilize their kidneys immediately after birth so that their kidney functions and therefore gene expression are reduced. In addition to difference in expressed genes, there is also a large variance in the numbers of expressed transcript isoforms per gene based on our Iso-seq data, ranging from 1.17 per gene in sciatic nerve to 3.4 per gene in hatchling brain (**Figure 4A**). This result unravels a highly dynamic pattern of transcription landscape among tissues in the western painted turtle.

**Figure 4:**
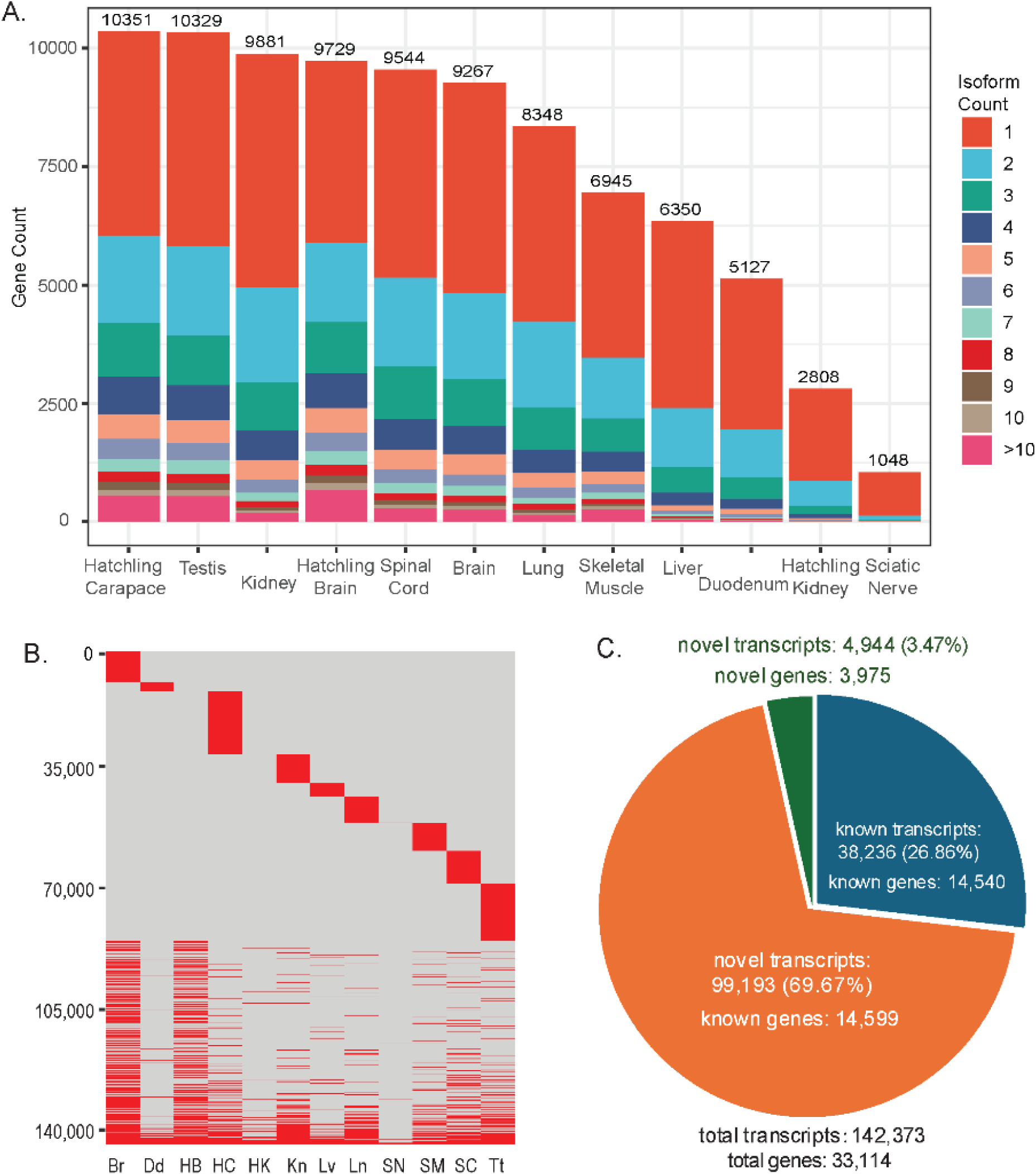
Characterization of transcription isoforms in 12 tissues of western painted turtle using PacBio Iso-Seq data. **A.** Distribution of detected genes and their transcript isoforms in the 12 tissues. **B.** Distribution of transcript isoforms across the 12 tissues. Each row represents a transcript isoform; numbers of isoforms are listed on the Y-axis. Each column represents a tissue (HC: Hatchling Carapace; Tt: Testis; Kn: Kidney; HB: Hatchling Brain; SC: Spinal Cord; Br: Brain; Ln: Lung; SM: Skeletal Muscle; Lv, Liver; Dd: Duodenum; HK: Hatchling Kidney; SN: Sciatic Nerve). For each tissue, red indicates the presence of a transcript isoform and light grey indicates the absence of the transcript isoform. **C.** Distribution of transcript isoforms detected by Iso-Seq. Most of transcript isoforms are not present in the NCBI genome annotation.

**Figure 5:**
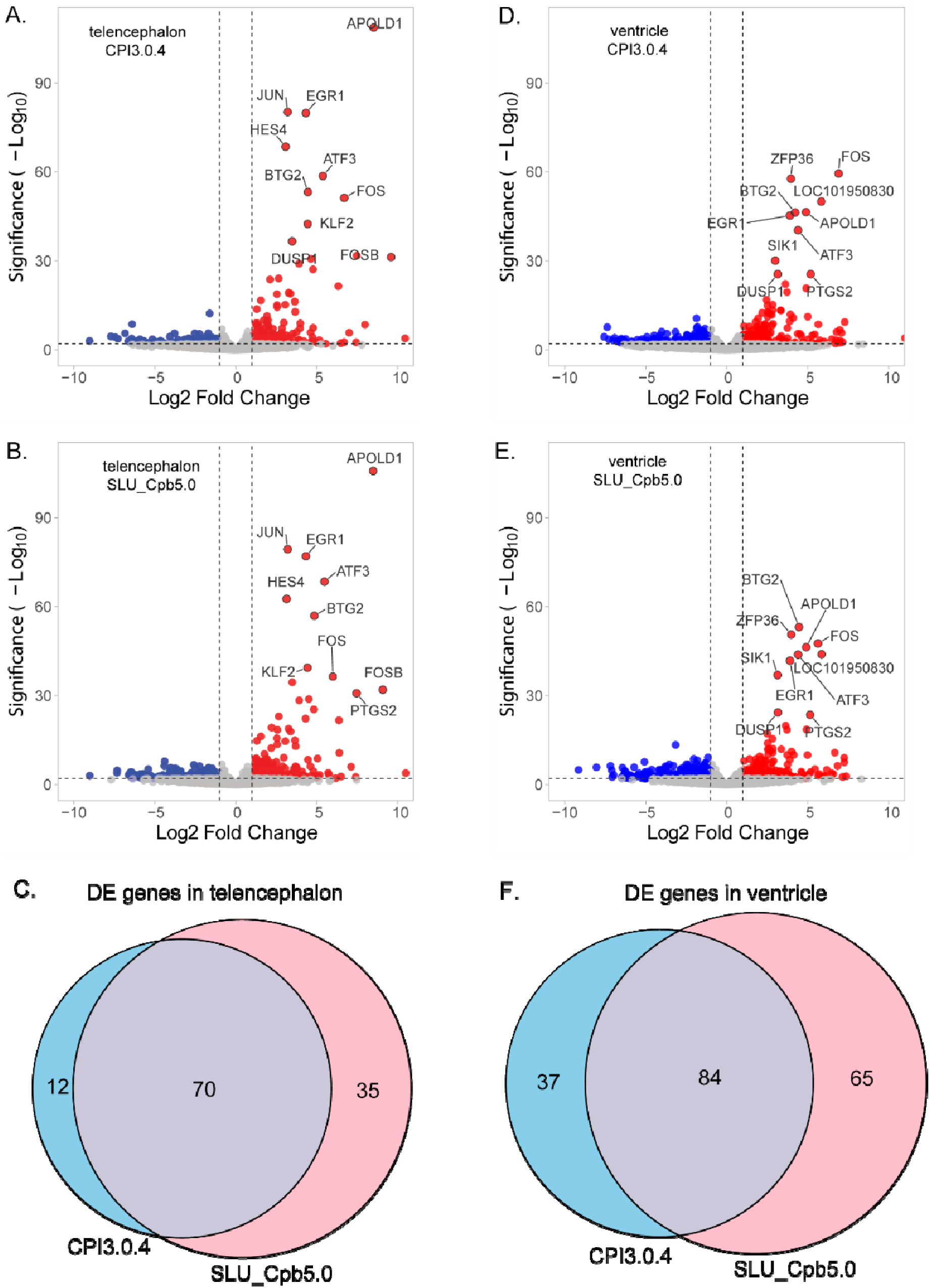
Impact on new genome assembly and annotation on differential gene expression analysis using RNA-seq data. **A**. Volcano plot showing differential gene expression analysis results from RNA-seq data in adult telencephalon using CPI3.0.4 assembly. **B**. Volcano plot showing differential gene expression analysis results from RNA-seq data in adult telencephalon using SLU_Cpb5.0 assembly. **C.** Venn diagram showing shared and assembly-specific DE genes in adult telencephalon. **D**. Volcano plot showing differential gene expression analysis results from RNA-seq data in adult ventricle using CPI3.0.4 assembly. Names of the top 10 DE genes are provided in each plot of A-D. **E, F**. Venn diagrams of differentially expressed genes in telencephalon and ventricle that were identified using different assemblies.

We also examined tissue-specific expression patterns of transcript isoforms (**Figure 4B**). Only 398 transcript isoforms are detected in all 12 tissues (**Figure 4B, Additional Table S11**), which are likely housekeeping transcript isoforms. It is noted that the number of transcript isoforms present in all tissues depends on the types and numbers of tissues examined, as well as sequencing depth. Thus, the number of shared transcript isoforms may differ if other tissues are examined or sequencing depth changes. Gene Ontology (GO) analysis suggests that these shared transcripts are mainly involved in cell adhesion (cadherin binding), catalytic activity (ATP hydrolysis, GTPase), and translational regulation (**Additional Figure S9A**). Interestingly, similar GO groups are enriched in tissue specific transcription isoforms (**Additional Figure S9B-D**). These results suggest that genes of these functional categories tend to express multiple transcript isoforms with two patterns: some are shared by all tissues, while many are only expressed in specific tissues. Elucidating the different functional roles of shared and tissue-specific isoforms generated by the same gene is expected to provide new insights into the complexity of gene regulation.

By merging all transcript isoforms identified in each tissue, we identified a total number of 142,373 transcript isoforms. The vast majority of identified transcript isoforms (96.54%) belong to genes annotated by NCBI, or known genes (**Figure 4C**). However, only 39,830 (27.8%) have identical splicing junctions with NCBI annotated transcripts, while 69% (98,249) are novel transcripts of these known genes. In addition, we identified 4,970 transcript isoforms belonging to 4,325 novel genes, which were absent in NCBI annotation (**Additional Table S11**). As only non-coding novel transcripts have been filtered, all these novel genes contain an open reading frame (ORF). The full-length transcriptome identified by Iso-Seq data significantly improves the completeness of genome annotation. The genome annotation data with novel transcripts and genes identified using our Iso-Seq data in gtf format is available for download at the GitHub site for this project (https://github.com/JohnnyChen1113/SLU_Cpb5.0).

### New Hypoxia/Anoxia-related transcripts

An important purpose for resequencing and reannotating the genome of the western painted turtle was to enable deeper insights into the transcriptomic basis for anoxia tolerance in this unique species. As described below, we have reanalyzed the RNA-seq dataset from our previous study of anoxic stress in telencephalon and ventricle of western painted turtle that were submerged in anoxic water for 12 hours [33]. At the time of the original study, the differential expression analysis of these RNA-seq datasets used assembly CPI3.0.1 and outdated informatic tools, which enabled the identification of 19 differentially expressed genes from adult telencephalon, all of which increased and 27 differentially expressed genes in ventricle, 23 of which were upregulated and 4 downregulated [33]. Since that time, the *C. picta bellii* genome underwent considerable improvements such that a reanalysis of these original RNA-seq data using CPI3.0.4 should yield many more differentially expressed genes. Our aim was to demonstrate how our new assembly, SLU_Cpb5.0, created with PacBio HiFi reads and scaffolded using Hi-C data, would reveal new differentially expressed genes not observed with any previous assembly version.

To this end, we reanalyzed these 16 RNA-seq datasets (NCBI SRA: SRS385156-SRS385171) by using genome assemblies SLU_Cpb5.0 and CPI3.0.4 as reference. Differential expression analyses using DESeq2 [66] were performed for mapped RNA-seq reads in both assemblies. Because there was a large difference in the number of annotated gene features between the two assembles (26,230 for CPI3.0.4 vs 34,544 for SLU_Cpb5.0), we selected a raw p-value significance cut-off based on the adjust p-values for the DE analysis conducted with the new assembly, and applied it to the p-values of the DE analysis data derived from the CPI3.0.4 assembly. This was a conservative approach because it led to the inclusion of many genes from the old assembly analysis whose adjusted p-values were not significant.

For adult telencephalon, 165 genes were found to be significantly differentially expressed during anoxia with SLU_Cpb5.0, 63 (38%) of which were not significant with the analysis using the CPI3.0.4 (**Additional Tables S12-14**). The expression of slightly more than half of these (33) was unquantifiable using the CPI3.0.4 assembly. These novel genes were mostly protein coding (38; 60%), 15 of which had not been quantified previously. The remaining were novel long non-coding RNAs (24; 38%), most of which (18) had never been quantified before. A novel differentially expressed small non-coding RNA was discovered when these data were reanalyzed with SLU_Cpb5.0.

For adult ventricle, 270 genes were significantly differentially expressed during anoxia with SLU_Cpb5.0, 109 (40%) of which were novel to the new assembly analysis. Even more (66; 61%) could not be quantified with CPI3.0.4. More than 60% of these novel genes were protein coding (also 66), 28 of which had not been previously quantified. The remaining novel genes were mostly long non-coding RNAs (42; 39%), most of which (38) had never been quantified before. The same novel differentially expressed small non-coding RNA detected in telencephalon was also differentially expressed in ventricle. Taken together, this analysis shows a significant improvement in the sensitivity of SLU_Cpb5.0 for detecting changes in differential gene expression in the tissues of anoxic turtles.

## DISCUSSION

We have generated a new *C. picta bellii* reference genome, SLU_Cpb5.0, that possesses several improvements over the previous reference, CPI3.0.4. First, our new assembly was generated using four datasets, 10x Genomics pseudo-longreads, PacBio HiFi, Hi-C, and BioNano optical reads, that were combined to produce a chromosome-level assembly with far greater contiguity. A comparison of genome assemblies derived from short- and long-reads demonstrates that the short-read assemblies miss up to 11% of the genome [67]. Second, our new assembly was assembled from datasets that were obtained from the same individual, eliminating the possibility of errors introduced by sequence variations between individuals. Third, our new assembly was annotated using full-length or near full length transcripts from twelve tissues, as well as published RNAseq datasets, providing extensive empirical evidence for annotation. The result of these efforts is the one of the best assembled and annotated turtle genomes to date, and an undeniable improvement over the previous *C. picta bellii* reference assembly and annotation.

The ability to affordably sequence genomes from any species is rapidly advancing systematics and conservation biology, as well as opening new avenues of research into the genetic basis for commercially or medically important traits. The Vertebrate Genomes Project (VGP; [31]) aims to sequence the genome of every vertebrate species, and has established methods and standards for sequencing and assembling these genomes. For non-model organisms, reference genomes can be used to answer questions across different time scales, ranging from phylogenomics to physiology, as well as to elucidate mechanisms of evolution, and to support conservation efforts (reviewed in [68, 69]). Our near-complete genome will facilitate several lines of research. First, high quality genome assemblies allow improved annotation [31] and permit more complete identification of gene families. Turtles have expanded families of water-soluble and aromatic olfactory receptors [28, 70], and future characterization of these genes in painted turtles will improve our understanding of the chemical cues that they can detect, thereby informing their behavior in the wild. Second, turtles are a particularly endangered taxon [71]; our improved genome assembly should aid conservation genomics efforts for other turtles: It can be used to identify SNPs in distantly related species [72], and provide a blueprint to map annotations onto other turtle genomes using programs such as Liftoff [73]. Third, our new chromosome-level genome provides a basis for further studies on painted turtle physiology and evolution. For example, we have used our extensively annotated assembly to identify new anoxia-related genes that were missed using the CPI3.0.4 assembly. Fourth, our western painted turtle genome includes phasing information, ie variants are linked. Thus, our assembly will facilitate future haplotype-based population genetics (eg [74]), thereby permitting detection of chromosomal variants and allele-specific expression that may involve physiologically relevant genes under selection. Finally, our genome assembly will permit identification of selective sweeps, which may include loci that are critical for cold adaptation or anoxia tolerance.

The turtle whose genome we sequenced is from the center of the western painted turtle range, whereas the previous assembly utilized a turtle from central Washington state, at the western edge of the species range. This new genome will help in the establishment of essential biodiversity variables, such as genetic diversity, genetic differentiation, inbreeding, and effective population size, which are important for conservation planning [75], particularly of western *C.picta bellii* populations [49]. Furthermore, reference bias in phylogenetic studies, originating from the assembly methods used to generate the reference, and/or the origin of the individual from which the reference genome was sequenced, can compromise identification of regions under selection [76]. These problems will be reduced by utilizing our new assembly.

### Painted turtle mitochondrial variation

Our analysis of the *C.picta bellii* mitochondrial genome provides several insights of potential importance. First, western painted turtles are a species of conservation interest in southwestern Canada, even though populations of the species as a whole are stable [49, 77]. The polymorphisms that we have identified that are outside of the Control Region will assist in clarifying western painted turtle taxonomy, and in preserving painted turtle biodiversity. The minimal number of differences between our mitochondrial genome and that from the Korean *C. picta bellii* individual suggests that the invasive Korean population is derived from Midwest *C. picta bellii*, not from northwestern *C. picta bellii*; therefore the Korean *Chrysemys* turtles are unlikely to retain genetic variation of conservation interest. Second, our estimates of the *C. picta*-*C. dorsalis* split at 1-2Mya, and the Minnesota and Washington *C. picta belli* divergence at 300,000-600,000 years ago are somewhat older than, but not inconsistent with, those from consensus phylogenetic trees that were generated from morphological characters and constrained by molecular data [78]. The turtle mitochondrial mutation rates [59] are based on single mitochondrial genes rather than entire mitochondrial genomes; this difference could result in artificially low mutation rates, thereby increasing our divergence time estimates. The numerous assumptions and uncertainties underlying our mitochondrial genome-based time-since-last-common-ancestor estimates underscore the need for similar estimates using whole genomes. Nevertheless, they suggest that the western *C. picta bellii* populations diverged earlier than currently assumed, prior to the last glacial maximum, highlighting the need to conserve their biodiversity, and providing more time for this widespread species to have established its current range.

Mitochondrial VNTRs are widespread among mitochondrial genomes [61]. For example, domestic cats possess 3-14 80-82bp repeats [79]. The widespread presence of Control Region VNTRs among mitochondrial genomes –that are constrained for size– suggest that VNTRs could be functional. The HMG-box protein TFAM not only controls mitochondrial transcription, but also binds mt-DNA non-specifically, compacting it into nucleoids [80]. We observe the TFAM binding site: GN_10_G (Reverse complement: CN_10_C; [81, 82]) in VNTR1B, suggesting that these repeats may influence mt-DNA compaction. Alternatively, VNTRs may function to relieve torsional stress from mt-DNA transcription or replication by permitting alternate structures such as Z-DNA or theta structures [83]. The effects, if any, of variation in these regions is unknown, but they represent a previously uncharacterized source of variability between *Chrysemys* populations.

Some of the differences between the *Chrysemys* mitochondrial genomes that we have observed could result from assembly errors. Mitochondrial genomes that have been assembled from short-read sequences tend to delete repeats, but they are also highly accurate for SNPs [61]. Our mitochondrial genome was generated from long reads, but the other four *Chrysemys* mitochondrial genomes were generated using short read sequences [28, 52, 54]. Thus, the fewer number of VNTR1A and VNTR1B repeats in the Korean *C.picta* mitochondrial genomes could be artifactual, but assembly errors are unlikely to account for the expanded VNTR1A and VNTRR1B repeats in *C. dorsalis*, or for the variation within the VNTR1 repeats. Additionally, nuclear mitochondrial DNA segments (NUMTs; reviewed in [84]), could confound our analysis. To assess this possibility, we conducted an NCBI BLAST search of our reference genome GCF_011386835.1 using our mitochondrial genome CM062378 as the query. We identified seven potential NUMTs, the largest of which, on Chromosome 1, covered 4974bp and had 96% identity to the mitochondrial genome. These fragmented, diverged NUMTs are unlikely to impact our mitochondrial genome alignments.

### Transcriptomics

A major feature of our painted turtle genome is its annotation using full-length, or near full length transcript sequences that were generated using PacBio Iso-Seq data generated from the RNAs of 12 tissues. Extensive variation in transcript start sites, splice sites, and polyadenylation are identified by long-read sequencing, as exemplified by the ENCODE4 RNAseq datasets from mouse and human tissues [85]. Splicing can be coupled to transcription [86]; potentially this coupling can be studied in painted turtle liver, from which we have both PacBio HiFi genomic methylation data and Isoseq data. For rainbow trout, a non-model species, long read transcripts improved annotation of its genome [87]; our Iso-Seq datasets will help extend the characterization of transcript variation to turtles. Alternative transcripts in turtles can have important physiological functions eg [88]. Notably, Long-read isoform sequencing reveals tissue-specific isoform expression between active and hibernating brown bears (Ursus arctos; [89]. Alternative CDC20 isoforms control the duration of mitotic arrest [90]; a similar mechanism could regulate mitotic arrest during hibernation. Lastly, our annotation will facilitate the use of spatial transcriptomics technologies such as 10x Genomics Visium to address new questions concerning transcript expression in specific turtle tissues and cell types.

### Anoxia tolerance

The Western painted turtle is the most anoxia-tolerant vertebrate known, capable of surviving for more than 30 hours at room temperature [12] and 170 days at 3°C [5, 7]. Because stroke, myocardial ischemia, and cardiac arrest produce tissue damage resulting from oxygen deprivation, understanding why some vertebrate species, such as turtles, are better than others at living without oxygen is medically important [91]. A well-annotated, chromosome-level western painted turtle genome should facilitate comparative genomics approaches to identify genes, genetic regulatory pathways, and physiological processes that enable extreme anoxia tolerance. To demonstrate the utility of this new genome (SLU_Cpb5.0) in elucidating the mechanisms of anoxia tolerance, we reanalyzed the short-read data from the first RNA-seq study of anoxia from our group [33], which involved Illumina Hi-seq paired-end sequencing of mRNAs from the telencephalon and cardiac ventricle of adult painted turtles subjected to 24 hours of anoxia at 19°C. We have demonstrated that our further refinement of the painted turtle genome greatly improved our ability to detect differentially expressed transcripts in both tissues. Specifically, this amounted to increases in the number of detectable differentially expressed genes of 38% and 40% in telencephalon and cardiac ventricle, respectively. This result lends support to the notion that much more can potentially be learned by the continued refinements, resequencing, and reannotation of genomes from organisms that possess extreme physiological phenotypes, such as the western painted turtle.

The purpose of the anoxia data reanalysis was to demonstrate the utility of the revised assembly for new discovery; here we are providing only preliminary findings of a few noteworthy and novel differentially expressed genes. First, the analysis of the RNA-seq analysis with the old assembly (CPI3.0.4) did not detect any changes in the expression of oxygen-sensitive prolyl-hydroxylases, which are broadly thought be involved in adaptive responses of cells to hypoxia. However, our re-analysis SLU_Cpb5.0 revealed significant increases in a highly expressed gene in the ventricle, *Egln3* (egl-9 family hypoxia inducible factor 3), also known as prolyl hydroxylase domain-containing 3 (PHD3). PHD3 hydroxylates HIF-1a and HIF-2b; it is a negative regulator of HIF pathway(s), which is generally thought to be adaptive. This leads to the novel hypothesis that avoiding overactivation of HIF-dependent pathways may be a protective mechanism in the heart [92–94].

Another notable and novel change in the anoxic ventricle was the increase in TCDD inducible poly (ADP-ribose) polymerase (*Tiparp*), often referred to as PARP7, which is constitutively expressed at relatively high levels. PARPs are poly-ADP-ribosyltranferases, which transfer the ADP-ribose moiety from NAD+ onto target proteins [95]. PARP7 functions as an inhibitor of the interferon response to cytoplasmic nucleic acids. Recent work has characterized PARP7 as a cysteine-specific mono-ADP-ribosyltransferase that targets dozens of nuclear proteins, including TBK1 and NF-kB that indue proinflammatory cytokines, as well as FRA1, an AP-1 transcription factor whose stabilization by PARP7 ADP-ribosylation represses pro-apoptotic pathways [96, 97]. Thus, upregulation of PARP7 during anoxia may enhance survival by preventing anoxia-induced apoptosis.

The urotensin-II receptor (*Uts2r*, GPR14) was found to be significantly elevated during anoxia in the heart. Urotensin II is a highly conserved peptide first discovered in fish (Pearson et al 1980) that has strong vasoactive effects when it binds GPR14 (Ross et al 2010). In mammalian heart, GPR14 is localized to cardiac myocytes with little expression in the vasculature [98]. Urotensin II increases contractile force of rat ventricular papillary muscles [98] and activates sarcolemmal Na+/H+ exchange in ventricular myocytes [99]. Both responses would be adaptive during anoxia by defending contractility against the negative ionotropic effects of anoxia-induced lactic acidosis.

The expression levels of *Txnip*, which is constitutively expressed at high levels, was also further increased during anoxia. This gene encodes Thioredoxin Interacting Protein, which binds and inhibits the activity of thioredoxin [100]. Theoretically, the inhibition of thioredoxin should increase cellular ROS production, but because anoxic turtle heart avoids ROS production altogether [22, 101], the role of TXNIP may serve functions other than redox balance. Targeted deletion of *Txnip* in mammalian heart showed no effect on thioredoxin or on ROS but, instead, stimulated rates of glucose uptake without changes in GLUT1 and GLUT4 expression, which implies a higher rate of glucose utilization [102]. Therefore, the role of TXNIP may be more related to adjusting the balance of glycolytic and oxidative pathways of energy production toward anaerobic glycolysis.

No prior mRNAseq studies of anoxic turtles showed any changes in the expression level of genes directly associated with cellular excitability or excitation contraction coupling [33, 34]. However, our reanalysis of the earlier of these studies with SLU_Cpb5.0 showed a significant decrease in the expression of a gene orthologous to mammalian TCAF2, which is a protein that promotes trafficking of TRPM8 to the membrane and negatively regulates it once it is there. TRPM8 is a cold- and menthol-sensitive calcium and sodium channel [103]. Inactivation of this channel during anoxia should decrease the excitability of cardiomyocytes, which would support cardiac functional suppression, and also defend intracellular calcium homeostasis.

Like ventricle, the majority of the novel DE genes in telencephalon were downregulated, but a few notable ones showed increases and could be related to regulation of the neuroepithelium and the blood-brain barrier. These include *Snai1*, which encodes Snail Family Transcriptional Repressor 1, a zinc-finger transcription factor that suppresses E-cadherin [104] and permeabilizes the blood-brain barrier [105]. *Cfap210*, which encodes Cilia and Flagella-Associated Protein 210, is a little-understood component of ciliary microtubules [106]. Cilia are important for motility and are hubs of signal transduction; *Cfap210* also was downregulated, further suggesting a suppression of neuroepithelial function during anoxia.

A potentially important adaptive response in telencephalon is the increase in *Cirbp* (sometimes *Cirp*); it encodes Cold-Inducible RNA-binding Protein, which was first shown to be induced in cancer cells after exposure to cold shock, UV radiation, hypoxia, and other cellular stresses [107]. It was later found to be upregulated during hypothermia in mammalian organotypic hippocampal slice cultures [108]. CIRBP regulates mRNA polyadenylation, stability and translation through binding on 3’-untranslated regions of target transcripts [109]. Several studies have implicated CIRBP as anti-apoptotic and potentially neuroprotective following ischemia due to its upregulation during mild therapeutic hypothermia [110–113]. Therefore, it is reasonable to hypothesize that CIRBP may play a neuroprotective role during and following anoxia in turtle brain.

### Conclusions

This new *C. picta bellii* genome assembly is a step towards elucidating the range of *C. picta* genetic variation, as well as potential selective signatures of adaptation to the environments that they inhabit. In addition to advancing *Chryemys* phylogenetics, the more complete annotation of our chromosome-level genome assembly allows better identification of differentially expressed genes. Genomic signatures of cold adaptation will lead to loci that encode the drivers of anoxia tolerance.

### Limitations of the study

Jarvis et al. [114] compared a variety of genome assembly pipelines, and found no single assembly approach that lacked errors in scaffold or contig errors. The sequencing technologies used here are imperfect; a k-mer analysis of a T2T assembly found sequences that were missing in both PacBio HiFi and Illumina reads [115]. We have attempted to correct as many errors as we can find, but we expect that some errors remain in our assembly. Although NCBI has designated our new assembly as the “reference” for *C. picta bellii* based on its improved contiguity and annotation, no single reference genome represents the inherent variability in natural populations [116].

## MATERIALS AND METHODS

### Sample collection and DNA extraction

All genomic DNA used in this study was obtained from the same male western painted turtle, *Chrysemys picta bellii* (notch ID R12L10; Biosample SAMN13293682 (See **Additional Table S1** for NCBI accession numbers), which was purchased from Niles Biologicals (Sacramento, CA), who procured it from McLeod County, MN in 2017. All procedures involving turtles were performed in the Warren lab at Saint Louis University using protocols approved by the Saint Louis University IACUC. Turtles were euthanized by phenobarbital injection into the subcarapacial sinus. Fresh whole blood for 10x Genomics sequencing was collected from turtle R12L10 by cardiac puncture; other tissues were harvested, rinsed in PBS, and snap frozen in aluminum freeze-clamps cooled in liquid nitrogen, before being transferred to dry ice and stored at -80°C. Frozen adult liver tissue was used for PacBio HiFi sequencing and for BioNano optical mapping. Frozen adult skeletal muscle was used for Hi-C sequencing.

### 10x Genomics sequencing

DNA extraction, library preparation, and sequencing were performed at the McDonnell Genome Institute, Washington University School of Medicine, St. Louis, MO. DNA was extracted from fresh, whole blood collected from heart, using Qiagen MagAttract, then used to prepare a library with 10x Chromium Genome v2, and sequenced to approximately 66X coverage on the 10x Genomics Chromium platform. This technology partitions high molecular weight genomic DNA fragments into droplets, where they are amplified with unique sequence identifiers (barcodes), to mark closely linked sequences prior to Illumina “short read” sequencing [35]. Approximately 150Gb of sequence was obtained on a NovaSeq S4 300XP (Sequence Read Archive #SRX7705481; see **Additional Table S1** for dataset accession numbers). The sequence was assembled using the de novo assembly program Supernova v2.1.1, and this genome assembly was submitted to NCBI, accession #GCA_011386835.1. Unassembled, barcoded 10x Genomics reads were used in subsequent assemblies.

### PacBio HiFi Sequencing

Pacific Biosystems (PacBio) HiFi genomic sequencing was performed by the DNA Sequencing Center at Brigham Young University, Provo, UT. Briefly, High molecular weight (HMW) DNA was isolated and cleaned from turtle liver using a Qiagen Genomic-tip kit and columns. Once isolated, the DNA was sheared on a Megaruptor (Diagenode/Hologic), AMPure cleaned, and made into a SMRTbell adapted library following PacBio manufacturer’s protocol. Library size selection was performed on a Blue Pippin (Sage Science, Beverly MA), collecting DNA fragments 10kb and larger. The size-selected library was prepared to run on a Sequel II using SMRTLink recommendations. Three flow cells were used, and their datasets were merged. The raw sequencing data has been submitted to NCBI SRA database (NCBI BioProject PRJNA589899; Sequence Read Archive #SRX22568648).

### Hi-C Sequencing

A Hi-C library was generated using DNA isolated from adult male liver tissue by Arima Genomics, Inc., Carlsbad, CA, using the Arima High Coverage Hi-C kit, part #A410110, following manufacturer’s protocols. DNA was crosslinked using the standard input protocol, and digested with a cocktail of four restriction enzymes. The Hi-C library was prepared using the Swift Biosciences Accel-NGS 2S Plus DNA Library kit, and sequenced on an Illumina NovaSeq (Sequence Read Archive #SRX22568649).

### BioNano Optical Mapping

Ultra-high molecular weight DNA was extracted from liver tissue, and fluorescently labeled using DLE-1. Labeling and imaging were performed by McDonnell Genome Institute at Washington University in St. Louis according to manufacturer’s protocol (Bionano Genomics, San Diego CA). DNA molecules greater than 150kb with a minimum of 9 labels were collected for Bionano labeling and loaded on the Saphyr chip for imaging. These data were assembled using Bionano Solve v.3.7 software (Bionano Genomics) with nonhaplotype_noES_DLE1_saphyr.xml parameters and placed to the assembled genome for validation.

### Genome assembly

To achieve optimal assembly results, we tested different tools at each stage of the genome assembly process (**Additional Figure S2**). We evaluated the assemblies using various metrics, including N50, total length, total sequence number, and BUSCO scores, to determine the most effective assembly method [43]. Initial *de novo* assemblies of the turtle genome were generated by different tools that provide haplotype-resolved assemblies, including including Hifiasm [36], NextDenovo [37], and Wtdbg2 [38]. Hifiasm, which used both PacBio HiFi data and Hi-C data, yielded three assemblies (primary, haplotype1 & 2) that had the best metrics in most categories; thus this assembly was selected for further scaffolding. A first-round scaffolding of the three assemblies was performed using 10x Genomics data with tigmint [39] and ARCS [39, 117]. A second round of scaffolding for each assembly was completed using Hi-C data by 3D-DNA [40]. This process involved using Hi-C data to order and orient scaffolds and to fill gaps between the scaffolds. Manual curation was performed using Bionano data with Bionano’s official toolkit (Solve & Access [118]) to further refine the assembly. The final *Chrysemys picta bellii* assembly, SLU_Cpb5.0, was deposited to NCBI BioProject PRJNA589899, and was adopted by NCBI as the reference assembly for western painted turtle (Genome assembly ASM1138683v2 and RefSeq assembly accession GCF_011386835.1; **Additional Table S1**).

Estimated genome size and degree of heterozygosity of western painted turtle R10L12 was inferred using HiFi reads and 10x Genomics Illumina reads. K-mer frequency files of the two types of sequencing data were obtained by Jellyfish [119] based on a k-mer size of 21 (k = 21). These files were used as input for GenomeScope 2.0 [41] to estimate the genome size and heterozygosity level.

### Comparisons of genome assembly and BAC clones

Only the BACs published in Badenhorst (2015) [29] were assigned NCBI Accession numbers, and therefore could be evaluated. These were queried on the NCBI BLAST site in January, 2024. These BAC sequences were used as queries for BLASTN searches against whole-genome shotgun contigs (wgs) of *Chrysemys picta bellii* (taxid:8478) on NCBI, optimized for highly similar sequences, typically with the following algorithm parameters: Max target sequences =10; Expect threshold=0.001; Word size=256, Match/Mismatch scores= 4,-5; Gap costs =Existence:12,Extension:8; Filters: Low complexity regions; Species-specific repeats for *Chrysemys picta bellii*; Mask for lookup table only.

### Assembly and comparative analysis of mitochondrial genome

We used MitoHiFi [50] to assemble the mitochondrial genome, with the primary assembly output from Hifiasm as input, the *C. picta bellii* mitochondrial genome NC_023890.1 in FASTA and GenBank formats as a template, and including the --mitos parameter in order to use MITOS2 [51] for annotation.

A BLASTn search for sequences that are homologous to the *C. picta bellii* VNTR1A was performed on the NCBI BLAST server using the following parameters: expect threshold 0.05; word size 7; match-mismatch reward-penalty: 1,-1; gap costs: 1,-2; no masking (see RID SR0K7A9X01N). A similar BLASTn search for sequences that are homologous to the *C. picta bellii* VNTR1B was performed on the NCBI BLAST server using the following parameters: expect threshold 0.05; word size 7; match-mismatch reward-penalty: 1,-1; gap costs: 0,2; no masking. We used MUSCLE on the EMBL server (https://www.ebi.ac.uk/Tools/msa/muscle/), and NCBI Clustal alignment tools to align mitochondrial genomes.

### Genome Annotation

Annotation of the primary assembly was completed using the NCBI eukaryotic genome annotation pipeline [62]. In brief, repetitive sequences of the primary assembly were first masked by WindowMasker [120]. Known RefSeq transcripts and Model RefSeq transcripts that pass the contamination screen were aligned locally to the genome using BLAST to identify the locations at which transcripts aligned. Global re-alignment of these RefSeq transcripts at these locations was performed with Splign to improve the accuracy of splice sites [121]. Evidence-based prediction approaches were conducted by Gnomon using protein and transcriptome alignments [62]. RNA-seq data obtained from western painted turtle was used to align to the masked genome with HISAT2 [44]. The unmasked RefSeq genome sequence file was utilized to identify small non-coding RNAs (sncRNAs). Transfer RNAs (tRNAs) were predicted using tRNAscan-SE, while ribosomal RNAs (rRNAs), small nucleolar RNAs (snoRNAs), and small nuclear RNAs (snRNAs) were annotated through the alignment of eukaryotic RFAM Hidden Markov Models (HMMs) against the genome using Infernal’s cmsearch tool [122–124]. The genome annotation files are available on the NCBI FTP server (see Annotation dataset, **Additional Table S1**). Annotation of transposable elements and repetitive sequences was conducted using the primary assembly by EarlGrey with default parameters [63].

### PacBio Iso-Seq RNA sequencing and characterization of tissue-specific transcript isoforms

To provide a more complete genome annotation and to identify tissue-specific transcript isoforms, we conducted PacBio Iso-Seq RNA sequencing using 12 tissues from adult western painted turtles: testis, hindbrain, liver, kidney, lung, skeletal muscle, spinal cord, sciatic nerve, duodenum, hatchling carapace, hatchling kidney, hatchling brain. Tissues were snap frozen in liquid N_2_ and stored at -80°C prior to RNA extraction. RNA extraction from these tissues, preparation of Iso-Seq libraries and sequencing using PacBio SMRT cell (Sequel II) were performed by the DNA Sequencing Center at Brigham Young University, Provo, UT.

Iso-Seq subreads data were first converted into circular consensus sequences (CCS) using the ccs software (https://ccs.how/) with a minimum read quality of 0.9 (--min-rq 0.9) to obtain more CCS reads. These CCS reads were then processed to extract full-length (FL) reads using lima (https://lima.how/) with the Iso-Seq processing mode (--isoseq and --peek-guess). To remove chimeric reads, these FL reads were then used to generate full-length non-chimeric (FLNC) reads with the “refine” function of isoseq3 (https://isoseq.how/). This procedure yielded separate FLNC read files for each of the 12 tissues (**Additional Table S9**).

All the 12 FLNC reads files were merged into a single file using the “merge” function of bamtools [125]. The merged FLNC read file was aligned to the reference genome using pbmm2 (https://github.com/PacificBiosciences/pbmm2) to generate a bam file. This bam file was processed to merge redundant transcript models into isoforms using the ‘collapse’ function of TAMA [126], generating a BED file that was then converted to GTF format using the TAMA format conversion script (tama_convert_bed_gtf_ensembl_no_cds.py).

To evaluate the quality of these isoform annotations, including isoform quality descriptors, junctions, and transcript ends, short-read RNA-seq data (**Additional Table S5**) were mapped to the reference genome using HISAT2 [44], and used as input for SQANTI3 [65]for splice junction assessment. Based on these quality control results, Iso-Seq annotated transcripts with low RNA-seq support were filtered out. Filtered annotation files from the 12 samples were then merged using the “Merge” function in TAMA to generate single annotation file [126]. To assign names to transcripts, the Iso-Seq annotation file was compared with the NCBI RefSeq annotation using homemade Python scripts (https://github.com/JohnnyChen1113/SLU_Cpb5.0).

### Evaluations of the impact of new genome assembly on differential expression analysis using RNA-seq data

We retrieved raw sequencing reads of RNA-seq data obtained in turtles under anoxia exposure [33, 34]. RNA-seq data were aligned to the existing assembly (v3.0.1) and this new assembly using HISAT2 [44]. Raw read count mapped to each annotated gene were obtained using featureCounts [127]. Differential expression analysis was performed using DESeq2 [66].

## DECLARATIONS

### Ethics approval and consent to participate

NOT APPLICABLE

### Consent for publication

Not APPLICABLE

### Availability of data and materials

All data is publicly available. Sequence data has been uploaded to NCBI, with accessions listed in Additional Table 2. A script that was used has been uploaded to GitHub.

### Author contributions

DEW, PM, and JRBJr conceived the study. DEW provided turtles, laboratory space and supplies. JC assembled genomes, performed annotations, and analyzed RNA-seq datasets, with input from JRBJr, DEW, and ZL. MK generated and analyzed BioNano optical maps. JRBJr wrote the initial manuscript draft; JRBJr, DEW, ZL and JC wrote sections, and revised the manuscript. JRBJr, DEW, and ZL provided funding. JC and JRBJr are equal contributors to this work and can reference this work as a first authorship paper in their curriculum vitae.

## Acknowledgements

Bob Fulton, Chad Tomlinson for 10x Genomics sequencing and assembly.

LaDeana Hillier for initial analysis of the 10x Genomics assembly.

Kaitlyn Bagwill for assistance in initial PacBio HiFi assembly.

Jim Henderson for initial BUSCO analysis.

Natasha Schneider, Nelson Membreno, and Michael/Mickey Arias for tissue dissections.

Paul Hime for advice on BAC-genome alignments

## REFERENCES

1. Valenzuela N: The painted turtle, Chrysemys picta: a model system for vertebrate evolution, ecology, and human health. Cold Spring Harb Protoc 2009, 2009:pdb emo124.

2. Cordero GA: Re-emergence of the Painted Turtle (Chrysemys picta) as a reference species for evo-devo. Evol Dev 2014, 16:184–188.

3. Moustakas-Verho JE, Cebra-Thomas J, Gilbert SF: Patterning of the turtle shell. Curr Opin Genet Dev 2017, 45:124–131.

4. Alibardi L: Development, structure, and protein composition of the corneous beak in turtles. Anat Rec (Hoboken) 2021, 304:2703–2725.

5. Ultsch GR, Jackson DC: Long-term submergence at 3 degrees C of the turtle Chrysemys picta bellii in normoxic and severely hypoxic water. III. Effects of changes in ambient PO2 and subsequent air breathing. J Exp Biol 1982, 97:87–99.

6. Jackson DC: Living without oxygen: lessons from the freshwater turtle. Comp Biochem Physiol A Mol Integr Physiol 2000, 125:299–315.

7. Odegard DT, Sonnenfelt MA, Bledsoe JG, Keenan SW, Hill CA, Warren DE: Changes in the material properties of the shell during simulated aquatic hibernation in the anoxia-tolerant painted turtle. J Exp Biol 2018, 221.

8. Storey KB, Storey JM, Brooks SP, Churchill TA, Brooks RJ: Hatchling turtles survive freezing during winter hibernation. Proc Natl Acad Sci U S A 1988, 85:8350–8354.

9. Paukstis GL, Shuman RD, Janzen FJ: Supercooling and freeze tolerance in hatchling painted turtles (Chrysemys picta). Canadian Journal of Zoology 1989, 67:1082–1084.

10. Packard GC, Packard MJ: Hatchling painted turtles (Chrysemys picta) survive exposure to subzero temperatures during hibernation by avoiding freezing. Journal of Comparative Physiology B 1993, 163:147–152.

11. Costanzo JP, Litzgus JD, Iverson JB, Lee RE: Seasonal changes in physiology and development of cold hardiness in the hatchling painted turtle Chrysemys picta. J Exp Biol 2000, 203:3459–3470.

12. Johlin JM, Moreland FB: Studies of the blood picture of the turtle after complete anoxia. J Biol Chem 1933, 103:107–114.

13. Starkey DE, Shaffer HB, Burke RL, Forstner MR, Iverson JB, Janzen FJ, Rhodin AG, Ultsch GR: Molecular systematics, phylogeography, and the effects of Pleistocene glaciation in the painted turtle (Chrysemys picta) complex. Evolution 2003, 57:119–128.

14. Seidel ME, Ernst CH: A Systematic Review of the Turtle Family Emydidae. Vertebrate Zoology 2017, 67:1–122.

15. Ernst CH, Lovich JE: Turtles of the United States and Canada. Second edn: The Johns Hopkins University Press; 2009.

16. Barela KL, Olson DH: Mapping the Western Pond Turtle (Actinemys marmorata) and Painted Turtle (Chrysemys picta) in western North America. Northwestern Naturalist 2014, 95:1–12.

17. Musacchia X: The viability of Chrysemys picta submerged at various temperatures. Physiological Zoology 1959, 32:47-50.

18. Wasser JS, Inman KC, Arendt EA, Lawler RG, Jackson DC: 31P-NMR measurements of pHi and high-energy phosphates in isolated turtle hearts during anoxia and acidosis. Am J Physiol 1990, 259:R521–530.

19. Jackson DC, Warburton SJ, Meinertz EA, Lawler RG, Wasser JS: The effect of prolonged anoxia at 3 degrees C on tissue high energy phosphates and phosphodiesters in turtles: a 31P-NMR study. J Comp Physiol [B] 1995, 165:77–84.

20. Bundgaard A, James AM, Gruszczyk AV, Martin J, Murphy MP, Fago A: Metabolic adaptations during extreme anoxia in the turtle heart and their implications for ischemia-reperfusion injury. Scientific reports 2019, 9:2850–2850.

21. Pamenter ME, Richards MD, Buck LT: Anoxia-induced changes in reactive oxygen species and cyclic nucleotides in the painted turtle. J Comp Physiol B 2007, 177:473–481.

22. Bundgaard A, Gruszczyk AV, Prag HA, Williams C, McIntyre A, Ruhr IM, James AM, Galli GLJ, Murphy MP, Fago A: Low production of mitochondrial reactive oxygen species after anoxia and reoxygenation in turtle hearts. J Exp Biol 2023, 226.

23. Lipton P: Ischemic cell death in brain neurons. Physiol Rev 1999, 79:1431–1568.

24. Hochachka PW, Lutz PL: Mechanism, origin, and evolution of anoxia tolerance in animals. Comp Biochem Physiol B Biochem Mol Biol 2001, 130:435–459.

25. Herbert CV, Jackson DC: Temperature effects on the responses to prolonged submergence in the turtle *Chrysemys picta* bellii. II. Metabolic rate, blood acid-base and ionic changes, and cardiovascular function in aerated and anoxic water. Physiol Zool 1985, 56:670–681.

26. Daw JC, Wenger DP, Berne RM: Relationship between cardiac glycogen and tolerance to anoxia in the western painted turtle, Chrysemys picta bellii. Comp Biochem Physiol 1967, 22:69–73.

27. Warren DE, Jackson DC: The metabolic consequences of repeated anoxic stress in the western painted turtle, Chrysemys picta bellii. Comp Biochem Physiol A Mol Integr Physiol 2017, 203:1–8.

28. Shaffer HB, Minx P, Warren DE, Shedlock AM, Thomson RC, Valenzuela N, Abramyan J, Amemiya CT, Badenhorst D, Biggar KK, et al: The western painted turtle genome, a model for the evolution of extreme physiological adaptations in a slowly evolving lineage. Genome Biol 2013, 14:R28.

29. Badenhorst D, Hillier LW, Literman R, Montiel EE, Radhakrishnan S, Shen Y, Minx P, Janes DE, Warren WC, Edwards SV, Valenzuela N: Physical Mapping and Refinement of the Painted Turtle Genome (Chrysemys picta) Inform Amniote Genome Evolution and Challenge Turtle-Bird Chromosomal Conservation. Genome Biol Evol 2015, 7:2038–2050.

30. Lee LS, Navarro-Dominguez BM, Wu Z, Montiel EE, Badenhorst D, Bista B, Gessler TB, Valenzuela N: Karyotypic Evolution of Sauropsid Vertebrates Illuminated by Optical and Physical Mapping of the Painted Turtle and Slider Turtle Genomes. Genes (Basel) 2020, 11.

31. Rhie A, McCarthy SA, Fedrigo O, Damas J, Formenti G, Koren S, Uliano-Silva M, Chow W, Fungtammasan A, Kim J, et al: Towards complete and error-free genome assemblies of all vertebrate species. Nature 2021, 592:737–746.

32. Leung SK, Jeffries AR, Castanho I, Jordan BT, Moore K, Davies JP, Dempster EL, Bray NJ, O’Neill P, Tseng E, et al: Full-length transcript sequencing of human and mouse cerebral cortex identifies widespread isoform diversity and alternative splicing. Cell Rep 2021, 37:110022.

33. Keenan SW, Hill CA, Kandoth C, Buck LT, Warren DE: Transcriptomic Responses of the Heart and Brain to Anoxia in the Western Painted Turtle. PLoS One 2015, 10:e0131669.

34. Fanter CE, Lin Z, Keenan SW, Janzen FJ, Mitchell TS, Warren DE: Development-specific transcriptomic profiling suggests new mechanisms for anoxic survival in the ventricle of overwintering turtles. J Exp Biol 2020, 223.

35. Zheng GX, Lau BT, Schnall-Levin M, Jarosz M, Bell JM, Hindson CM, Kyriazopoulou-Panagiotopoulou S, Masquelier DA, Merrill L, Terry JM, et al: Haplotyping germline and cancer genomes with high-throughput linked-read sequencing. Nat Biotechnol 2016, 34:303–311.

36. Cheng H, Concepcion GT, Feng X, Zhang H, Li H: Haplotype-resolved de novo assembly using phased assembly graphs with hifiasm. Nat Methods 2021, 18:170–175.

37. Hu J, Wang Z, Sun Z, Hu B, Ayoola AO, Liang F, Li J, Sandoval JR, Cooper DN, Ye K, et al: NextDenovo: an efficient error correction and accurate assembly tool for noisy long reads. Genome Biol 2024, 25:107.

38. Ruan J, Li H: Fast and accurate long-read assembly with wtdbg2. Nat Methods 2020, 17:155–158.

39. Jackman SD, Coombe L, Chu J, Warren RL, Vandervalk BP, Yeo S, Xue Z, Mohamadi H, Bohlmann J, Jones SJM, Birol I: Tigmint: correcting assembly errors using linked reads from large molecules. BMC Bioinformatics 2018, 19:393.

40. Dudchenko O, Batra SS, Omer AD, Nyquist SK, Hoeger M, Durand NC, Shamim MS, Machol I, Lander ES, Aiden AP, Aiden EL: De novo assembly of the Aedes aegypti genome using Hi-C yields chromosome-length scaffolds. Science 2017, 356:92–95.

41. Ranallo-Benavidez TR, Jaron KS, Schatz MC: GenomeScope 2.0 and Smudgeplot for reference-free profiling of polyploid genomes. Nat Commun 2020, 11:1432.

42. Ren Y, Zhang Q, Yan X, Hou D, Huang H, Li C, Rao D, Li Y: Genomic insights into the evolution of the critically endangered soft-shelled turtle Rafetus swinhoei. Mol Ecol Resour 2022, 22:1972–1985.

43. Manni M, Berkeley MR, Seppey M, Simao FA, Zdobnov EM: BUSCO Update: Novel and Streamlined Workflows along with Broader and Deeper Phylogenetic Coverage for Scoring of Eukaryotic, Prokaryotic, and Viral Genomes. Mol Biol Evol 2021, 38:4647–4654.

44. Kim D, Paggi JM, Park C, Bennett C, Salzberg SL: Graph-based genome alignment and genotyping with HISAT2 and HISAT-genotype. Nat Biotechnol 2019, 37:907–915.

45. Simison WB, Parham JF, Papenfuss TJ, Lam AW, Henderson JB: An Annotated Chromosome-Level Reference Genome of the Red-Eared Slider Turtle (Trachemys scripta elegans). Genome Biol Evol 2020, 12:456–462.

46. Li H, Durbin R: Genome assembly in the telomere-to-telomere era. Nat Rev Genet 2024, 25:658–670.

47. Bista B, Gonzalez-Rodelas L, Alvarez-Gonzalez L, Wu ZQ, Montiel EE, Lee LS, Badenhorst DB, Radhakrishnan S, Literman R, Navarro-Dominguez B, et al: De novo genome assemblies of two cryptodiran turtles with ZZ/ZW and XX/XY sex chromosomes provide insights into patterns of genome reshuffling and uncover novel 3D genome folding in amniotes. Genome Res 2024, 34:1553–1569.

48. Suomalainen A, Nunnari J: Mitochondria at the crossroads of health and disease. Cell 2024, 187:2601–2627.

49. Jensen EL, Govindarajulu P, Russello MA: When the shoe doesn’t fit: applying conservation unit concepts to western painted turtles at their northern periphery. Conservation Genetics 2014, 15:261–274.

50. Uliano-Silva M, Ferreira J, Krasheninnikova K, Darwin Tree of Life C, Formenti G, Abueg L, Torrance J, Myers EW, Durbin R, Blaxter M, McCarthy SA: MitoHiFi: a python pipeline for mitochondrial genome assembly from PacBio high fidelity reads. BMC Bioinformatics 2023, 24:288.

51. Donath A, Juhling F, Al-Arab M, Bernhart SH, Reinhardt F, Stadler PF, Middendorf M, Bernt M: Improved annotation of protein-coding genes boundaries in metazoan mitochondrial genomes. Nucleic Acids Res 2019, 47:10543–10552.

52. Ji YE, Park KH, Choi JH, Park J, Sung HC, Lee DH: Complete mitochondrial genome of the southern painted turtle (Chrysemys dorsalis, Testudines: Emydidae) in Korea. Mitochondrial DNA B Resour 2024, 9:70–74.

53. Jiang JJ, Xia EH, Gao CW, Gao LZ: The complete mitochondrial genome of western painted turtle, Chrysemys picta bellii (Chrysemys, Emydidae). Mitochondrial DNA A DNA Mapp Seq Anal 2016, 27:787–788.

54. Park J, Park SM, Choi JH, Sung HC, Lee DH: Complete mitochondrial genome of the western painted turtle (Chrysemys picta bellii, Testudines: Emydidae) in Korea. Mitochondrial DNA B Resour 2023, 8:1316–1319.

55. Reese SA, Stewart ER, Crocker CE, Jackson DC, Ultsch GR: Geographic variation of the physiological response to overwintering in the painted turtle (Chrysemys picta). Physiol Biochem Zool 2004, 77:619–630.

56. Green RE, Malaspinas AS, Krause J, Briggs AW, Johnson PL, Uhler C, Meyer M, Good JM, Maricic T, Stenzel U, et al: A complete Neandertal mitochondrial genome sequence determined by high-throughput sequencing. Cell 2008, 134:416–426.

57. Levinstein Hallak K, Rosset S: Dating ancient splits in phylogenetic trees, with application to the human-Neanderthal split. *BMC Genom Data* 2024, 25:4.

58. Soares P, Ermini L, Thomson N, Mormina M, Rito T, Rohl A, Salas A, Oppenheimer S, Macaulay V, Richards MB: Correcting for purifying selection: an improved human mitochondrial molecular clock. Am J Hum Genet 2009, 84:740–759.

59. Allio R, Donega S, Galtier N, Nabholz B: Large Variation in the Ratio of Mitochondrial to Nuclear Mutation Rate across Animals: Implications for Genetic Diversity and the Use of Mitochondrial DNA as a Molecular Marker. Mol Biol Evol 2017, 34:2762–2772.

60. Bernacki LE, Kilpatrick CW: Structural Variation of the Turtle Mitochondrial Control Region. J Mol Evol 2020, 88:618–640.

61. Formenti G, Rhie A, Balacco J, Haase B, Mountcastle J, Fedrigo O, Brown S, Capodiferro MR, Al-Ajli FO, Ambrosini R, et al: Complete vertebrate mitogenomes reveal widespread repeats and gene duplications. Genome Biol 2021, 22:120.

62. Thibaud-Nissen F, DiCuccio M, Hlavina W, Kimchi A, Kitts PA, Murphy TD, Pruitt KD, Souvorov A: P8008 The NCBI Eukaryotic Genome Annotation Pipeline. Journal of Animal Science 2016, 94:184–184.

63. Baril T, Galbraith J, Hayward A: Earl Grey: A Fully Automated User-Friendly Transposable Element Annotation and Analysis Pipeline. Mol Biol Evol 2024, 41.

64. Lee Y, Rio DC: Mechanisms and Regulation of Alternative Pre-mRNA Splicing. Annu Rev Biochem 2015, 84:291–323.

65. Pardo-Palacios FJ, Arzalluz-Luque A, Kondratova L, Salguero P, Mestre-Tomas J, Amorin R, Estevan-Morio E, Liu T, Nanni A, McIntyre L, et al: SQANTI3: curation of long-read transcriptomes for accurate identification of known and novel isoforms. Nat Methods 2024, 21:793–797.

66. Love MI, Huber W, Anders S: Moderated estimation of fold change and dispersion for RNA-seq data with DESeq2. Genome Biol 2014, 15:550.

67. Kim J, Lee C, Ko BJ, Yoo DA, Won S, Phillippy AM, Fedrigo O, Zhang G, Howe K, Wood J, et al: False gene and chromosome losses in genome assemblies caused by GC content variation and repeats. Genome Biol 2022, 23:204.

68. da Fonseca RR, Albrechtsen A, Themudo GE, Ramos-Madrigal J, Sibbesen JA, Maretty L, Zepeda-Mendoza ML, Campos PF, Heller R, Pereira RJ: Next-generation biology: Sequencing and data analysis approaches for non-model organisms. Marine Genomics 2016, 30:3–13.

69. Worley KC, Richards S, Rogers J: The value of new genome references. Exp Cell Res 2017, 358:433–438.

70. Bentley BP, Carrasco-Valenzuela T, Ramos EKS, Pawar H, Souza Arantes L, Alexander A, Banerjee SM, Masterson P, Kuhlwilm M, Pippel M, et al: Divergent sensory and immune gene evolution in sea turtles with contrasting demographic and life histories. Proc Natl Acad Sci U S A 2023, 120:e2201076120.

71. Gumbs R, Scott O, Bates R, Bohm M, Forest F, Gray CL, Hoffmann M, Kane D, Low C, Pearse WD, et al: Global conservation status of the jawed vertebrate Tree of Life. Nat Commun 2024, 15:1101.

72. Galla SJ, Forsdick NJ, Brown L, Hoeppner MP, Knapp M, Maloney RF, Moraga R, Santure AW, Steeves TE: Reference Genomes from Distantly Related Species Can Be Used for Discovery of Single Nucleotide Polymorphisms to Inform Conservation Management. Genes (Basel) 2019, 10:9.

73. Shumate A, Salzberg SL: Liftoff: accurate mapping of gene annotations. Bioinformatics 2021, 37:1639–1643.

74. Meier JI, Salazar PA, Kucka M, Davies RW, Dreau A, Aldas I, Box Power O, Nadeau NJ, Bridle JR, Rolian C, et al: Haplotype tagging reveals parallel formation of hybrid races in two butterfly species. Proc Natl Acad Sci U S A 2021, 118.

75. Hogg CJ: Translating genomic advances into biodiversity conservation. Nat Rev Genet 2024, 25:362–373.

76. Thorburn DJ, Sagonas K, Binzer-Panchal M, Chain FJJ, Feulner PGD, Bornberg-Bauer E, Reusch TBH, Samonte-Padilla IE, Milinski M, Lenz TL, Eizaguirre C: Origin matters: Using a local reference genome improves measures in population genomics. Mol Ecol Resour 2023, 23:1706–1723.

77. Jensen EL, Govindarajulu P, Russello MA: Genetic Assessment of Taxonomic Uncertainty in Painted Turtles. Journal of Herpetology 2015, 49:314–324.

78. Jasinski SE: A new species of Chrysemys (Emydidae: Deirochelyinae) from the latest Miocene-Early Pliocene of Tennessee, USA and its implications for the evolution of painted turtles. Zoological Journal of the Linnean Society 2023, 198:149–183.

79. Patterson EC, Lall GM, Neumann R, Ottolini B, Batini C, Sacchini F, Foster AP, Wetton JH, Jobling MA: Mitogenome sequences of domestic cats demonstrate lineage expansions and dynamic mutation processes in a mitochondrial minisatellite. BMC Genomics 2023, 24:690.

80. Menger KE, Rodriguez-Luis A, Chapman J, Nicholls TJ: Controlling the topology of mammalian mitochondrial DNA. Open Biol 2021, 11:210168.

81. Cuppari A, Fernandez-Millan P, Battistini F, Tarres-Sole A, Lyonnais S, Iruela G, Ruiz-Lopez E, Enciso Y, Rubio-Cosials A, Prohens R, et al: DNA specificities modulate the binding of human transcription factor A to mitochondrial DNA control region. Nucleic Acids Res 2019, 47:6519–6537.

82. Choi WS, Garcia-Diaz M: A minimal motif for sequence recognition by mitochondrial transcription factor A (TFAM). Nucleic Acids Res 2022, 50:322–332.

83. Higgins NP, Vologodskii AV: Topological Behavior of Plasmid DNA. Microbiol Spectr 2015, 3.

84. Xue L, Moreira JD, Smith KK, Fetterman JL: The Mighty NUMT: Mitochondrial DNA Flexing Its Code in the Nuclear Genome. Biomolecules 2023, 13.

85. Reese F, Williams B, Balderrama-Gutierrez G, Wyman D, Celik MH, Rebboah E, Rezaie N, Trout D, Razavi-Mohseni M, Jiang Y, et al: The ENCODE4 long-read RNA-seq collection reveals distinct classes of transcript structure diversity. bioRxiv 2023.

86. Shenasa H, Bentley DL: Pre-mRNA splicing and its cotranscriptional connections. Trends Genet 2023, 39:672–685.

87. Ali A, Thorgaard GH, Salem M: PacBio Iso-Seq Improves the Rainbow Trout Genome Annotation and Identifies Alternative Splicing Associated With Economically Important Phenotypes. Front Genet 2021, 12:683408.

88. Mizoguchi B, Valenzuela N: Alternative splicing and thermosensitive expression of Dmrt1 during urogenital development in the painted turtle, Chrysemys picta. PeerJ 2020, 8:e8639.

89. Tseng E, Underwood JG, Evans Hutzenbiler BD, Trojahn S, Kingham B, Shevchenko O, Bernberg E, Vierra M, Robbins CT, Jansen HT, Kelley JL: Long-read isoform sequencing reveals tissue-specific isoform expression between active and hibernating brown bears (Ursus arctos). G3 (Bethesda) 2022, 12.

90. Tsang MJ, Cheeseman IM: Alternative CDC20 translational isoforms tune mitotic arrest duration. Nature 2023, 617:154–161.

91. Bickler PE: Clinical perspectives: neuroprotection lessons from hypoxia-tolerant organisms. J Exp Biol 2004, 207:3243–3249.

92. Henze AT, Riedel J, Diem T, Wenner J, Flamme I, Pouyseggur J, Plate KH, Acker T: Prolyl hydroxylases 2 and 3 act in gliomas as protective negative feedback regulators of hypoxia-inducible factors. Cancer Res 2010, 70:357–366.

93. Deng H, Wang Z, Zhu C, Chen Z: Prolyl hydroxylase domain enzyme PHD2 inhibits proliferation and metabolism in non-small cell lung cancer cells in HIF-dependent and HIF-independent manners. Front Oncol 2024, 14:1370393.

94. Kozlova N, Wottawa M, Katschinski DM, Kristiansen G, Kietzmann T: Hypoxia-inducible factor prolyl hydroxylase 2 (PHD2) is a direct regulator of epidermal growth factor receptor (EGFR) signaling in breast cancer. Oncotarget 2017, 8:9885–9898.

95. Bock FJ, Todorova TT, Chang P: RNA Regulation by Poly(ADP-Ribose) Polymerases. Mol Cell 2015, 58:959–969.

96. Manetsch P, Bohi F, Nowak K, Leslie Pedrioli DM, Hottiger MO: PARP7-mediated ADP-ribosylation of FRA1 promotes cancer cell growth by repressing IRF1- and IRF3-dependent apoptosis. Proc Natl Acad Sci U S A 2023, 120:e2309047120.

97. Manetsch P, Hottiger MO: Unleashing viral mimicry: A combinatorial strategy to enhance the efficacy of PARP7 inhibitors. Bioessays 2025, 47:e2400087.

98. Gong H, Wang YX, Zhu YZ, Wang WW, Wang MJ, Yao T, Zhu YC: Cellular distribution of GPR14 and the positive inotropic role of urotensin II in the myocardium in adult rat. J Appl Physiol (1985) 2004, 97:2228–2235.

99. Kato K, Yasutake M, Jia D, Snabaitis AK, Avkiran M, Kusama Y, Takano T, Mizuno K: Urotensin II activates sarcolemmal Na+/H+ exchanger in adult rat ventricular myocytes. J Cardiovasc Pharmacol 2010, 55:191–197.

100. Wang BF, Yoshioka J: The Emerging Role of Thioredoxin-Interacting Protein in Myocardial Ischemia/Reperfusion Injury. J Cardiovasc Pharmacol Ther 2017, 22:219–229.

101. Ruhr IM, McCourty H, Bajjig A, Crossley DA, 2nd, Shiels HA, Galli GLJ: Developmental plasticity of cardiac anoxia-tolerance in juvenile common snapping turtles ( Chelydra serpentina). Proc Biol Sci 2019, 286:20191072.

102. Yoshioka J, Imahashi K, Gabel SA, Chutkow WA, Burds AA, Gannon J, Schulze PC, MacGillivray C, London RE, Murphy E, Lee RT: Targeted deletion of thioredoxin-interacting protein regulates cardiac dysfunction in response to pressure overload. Circ Res 2007, 101:1328–1338.

103. Gkika D, Lemonnier L, Shapovalov G, Gordienko D, Poux C, Bernardini M, Bokhobza A, Bidaux G, Degerny C, Verreman K, et al: TRP channel-associated factors are a novel protein family that regulates TRPM8 trafficking and activity. J Cell Biol 2015, 208:89–107.

104. Batlle E, Sancho E, Franci C, Dominguez D, Monfar M, Baulida J, Garcia De Herreros A: The transcription factor snail is a repressor of E-cadherin gene expression in epithelial tumour cells. Nat Cell Biol 2000, 2:84–89.

105. Kim BJ, Hancock BM, Bermudez A, Del Cid N, Reyes E, van Sorge NM, Lauth X, Smurthwaite CA, Hilton BJ, Stotland A, et al: Bacterial induction of Snail1 contributes to blood-brain barrier disruption. J Clin Invest 2015, 125:2473–2483.

106. Gui M, Croft JT, Zabeo D, Acharya V, Kollman JM, Burgoyne T, Hoog JL, Brown A: SPACA9 is a lumenal protein of human ciliary singlet and doublet microtubules. Proc Natl Acad Sci U S A 2022, 119:e2207605119.

107. Lleonart ME: A new generation of proto-oncogenes: cold-inducible RNA binding proteins. Biochim Biophys Acta 2010, 1805:43–52.

108. Tong G, Endersfelder S, Rosenthal LM, Wollersheim S, Sauer IM, Buhrer C, Berger F, Schmitt KR: Effects of moderate and deep hypothermia on RNA-binding proteins RBM3 and CIRP expressions in murine hippocampal brain slices. Brain Res 2013, 1504:74–84.

109. Corre M, Lebreton A: Regulation of cold-inducible RNA-binding protein (CIRBP) in response to cellular stresses. Biochimie 2024, 217:3–9.

110. Wu L, Sun HL, Gao Y, Hui KL, Xu MM, Zhong H, Duan ML: Therapeutic Hypothermia Enhances Cold-Inducible RNA-Binding Protein Expression and Inhibits Mitochondrial Apoptosis in a Rat Model of Cardiac Arrest. Mol Neurobiol 2017, 54:2697–2705.

111. Saito K, Fukuda N, Matsumoto T, Iribe Y, Tsunemi A, Kazama T, Yoshida-Noro C, Hayashi N: Moderate low temperature preserves the stemness of neural stem cells and suppresses apoptosis of the cells via activation of the cold-inducible RNA binding protein. Brain Res 2010, 1358:20–29.

112. Li S, Zhang Z, Xue J, Liu A, Zhang H: Cold-inducible RNA binding protein inhibits H(2)O(2)-induced apoptosis in rat cortical neurons. Brain Res 2012, 1441:47–52.

113. Zhang HT, Xue JH, Zhang ZW, Kong HB, Liu AJ, Li SC, Xu DG: Cold-inducible RNA-binding protein inhibits neuron apoptosis through the suppression of mitochondrial apoptosis. Brain Res 2015, 1622:474–483.

114. Jarvis ED, Formenti G, Rhie A, Guarracino A, Yang C, Wood J, Tracey A, Thibaud-Nissen F, Vollger MR, Porubsky D, et al: Semi-automated assembly of high-quality diploid human reference genomes. Nature 2022, 611:519–531.

115. Mc Cartney AM, Shafin K, Alonge M, Bzikadze AV, Formenti G, Fungtammasan A, Howe K, Jain C, Koren S, Logsdon GA, et al: Chasing perfection: validation and polishing strategies for telomere-to-telomere genome assemblies. Nat Methods 2022, 19:687–695.

116. Eisenstein M: Every base everywhere all at once: pangenomics comes of age. Nature 2023, 616:618–620.

117. Yeo S, Coombe L, Warren RL, Chu J, Birol I: ARCS: scaffolding genome drafts with linked reads. Bioinformatics 2018, 34:725–731.

118. Shelton JM, Coleman MC, Herndon N, Lu N, Lam ET, Anantharaman T, Sheth P, Brown SJ: Tools and pipelines for BioNano data: molecule assembly pipeline and FASTA super scaffolding tool. BMC Genomics 2015, 16:734.

119. Marçais G, Kingsford C: A fast, lock-free approach for efficient parallel counting of occurrences of k-mers. Bioinformatics 2011, 27:764–770.

120. Morgulis A, Gertz EM, Schaffer AA, Agarwala R: WindowMasker: window-based masker for sequenced genomes. Bioinformatics 2006, 22:134–141.

121. Kapustin Y, Souvorov A, Tatusova T, Lipman D: Splign: algorithms for computing spliced alignments with identification of paralogs. Biol Direct 2008, 3:20.

122. Nawrocki EP, Eddy SR: Infernal 1.1: 100-fold faster RNA homology searches. Bioinformatics 2013, 29:2933–2935.

123. Kalvari I, Argasinska J, Quinones-Olvera N, Nawrocki EP, Rivas E, Eddy SR, Bateman A, Finn RD, Petrov AI: Rfam 13.0: shifting to a genome-centric resource for non-coding RNA families. Nucleic Acids Res 2018, 46:D335–D342.

124. Chan PP, Lin BY, Mak AJ, Lowe TM: tRNAscan-SE 2.0: improved detection and functional classification of transfer RNA genes. Nucleic Acids Res 2021, 49:9077–9096.

125. Barnett DW, Garrison EK, Quinlan AR, Stromberg MP, Marth GT: BamTools: a C++ API and toolkit for analyzing and managing BAM files. Bioinformatics 2011, 27:1691–1692.

126. Kuo RI, Cheng Y, Zhang R, Brown JWS, Smith J, Archibald AL, Burt DW: Illuminating the dark side of the human transcriptome with long read transcript sequencing. BMC Genomics 2020, 21:751.

127. Liao Y, Smyth GK, Shi W: featureCounts: an efficient general purpose program for assigning sequence reads to genomic features. Bioinformatics 2014, 30:923–930.

128. Krzywinski M, Schein J, Birol I, Connors J, Gascoyne R, Horsman D, Jones SJ, Marra MA: Circos: an information aesthetic for comparative genomics. Genome Res 2009, 19:1639–1645.

129. He W, Yang J, Jing Y, Xu L, Yu K, Fang X: NGenomeSyn: an easy-to-use and flexible tool for publication-ready visualization of syntenic relationships across multiple genomes. Bioinformatics 2023, 39.

