## Supplemental Figures for "A new chromosome-level genome assembly for western painted turtle *Chrysemys picta belli*, a model for extreme physiological adaptations"

**Additional Figures:**

**•	Additional Figure S1: Geological distribution of western painted turtle in North America.**

**•	Additional Figure S2: Flowchart describing procedures of genome assembly in this study.**

**•	Additional Figure S3: Read length and quality distribution of PacBio HiFi sequencing data.**

**•	Additional Figure S4: Hi-C contact heatmap visualizing chromosomal interactions. The heatmaps display interaction patterns for three assemblies.**

**•	Additional Figure S5: Examples of manual corrections in chromosome assemblies using Hi-C contact maps and BioNano optical mapping**

**•	Additional Figure S6: The k-mer distribution of different sequencing reads of turtle genomic DNA.**

**•	Additional Figure S7: Graphic displays of BLAST localization of representative BACs to *C.picta bellii* genome assemblies.**

**•	Additional Figure S8: Mitochondrial VNTR1 repeats.**

**•	Additional Figure S9: Gene Ontology (GO) analyses reveal enriched biological functions of shared and tissue specific transcript isoforms**

**Additional Table Legends**

**Additional References**

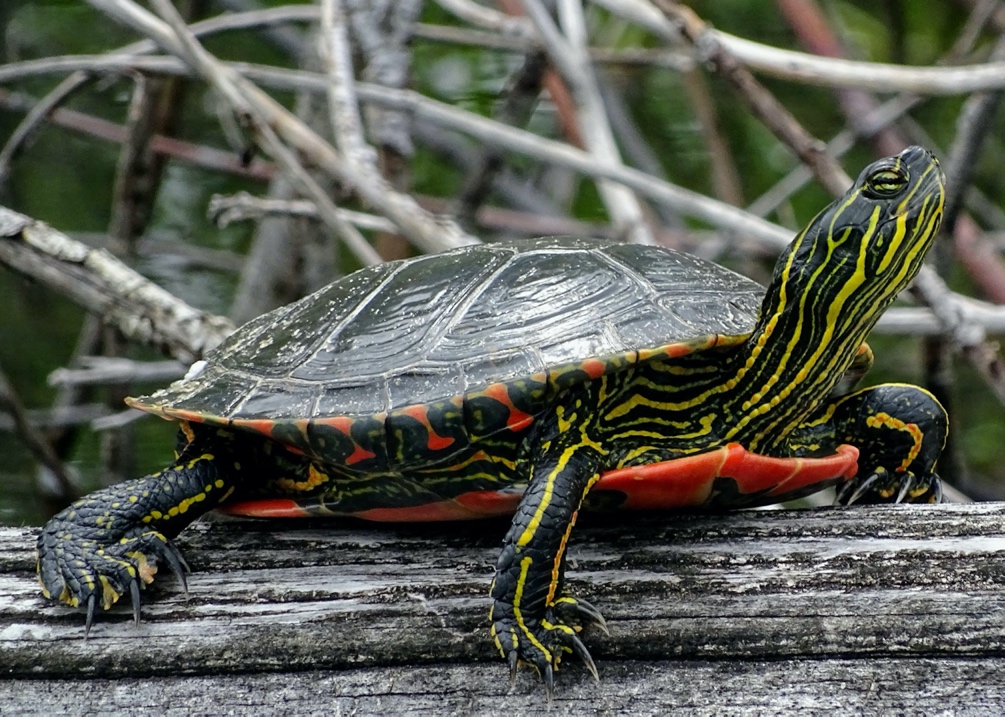

**A**

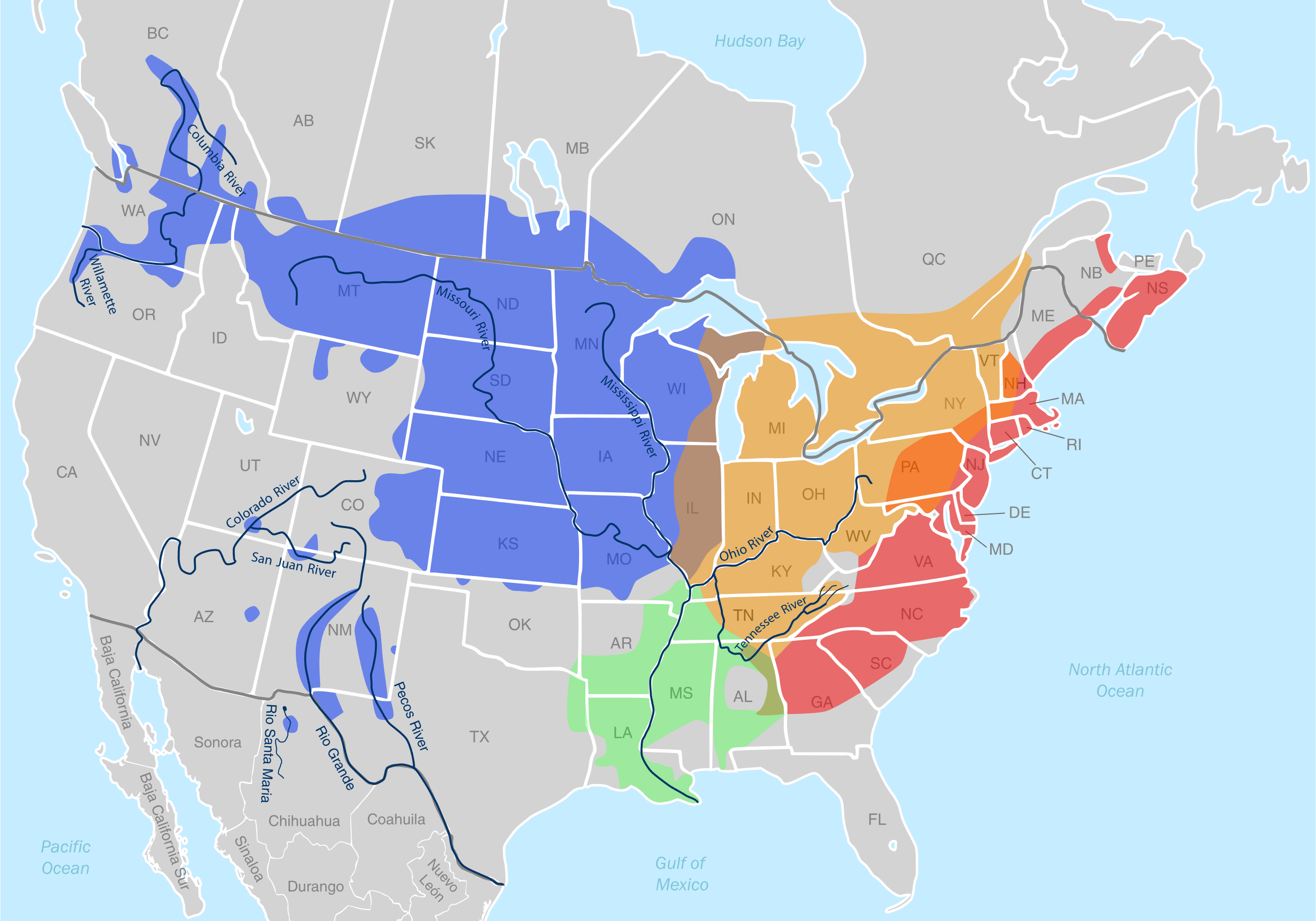

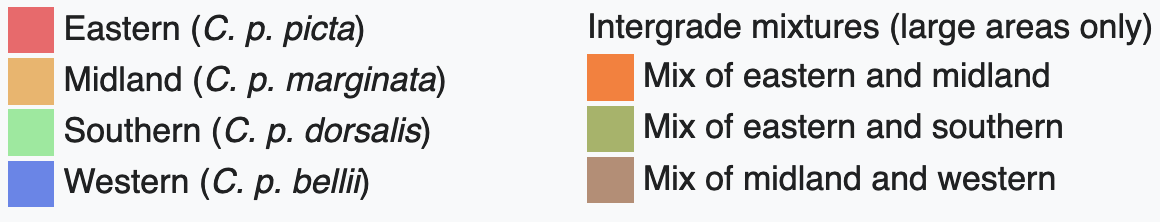

McLeod County, MN

**B**

**B**

**_(9(99(_**

**Legend to Additional Figure S1: Western Painted turtle, *Chrysemys picta bellii*.**

(**A**) A typical Western painted turtle. Photograph from iStock/Dave Acheson photography. (**B**) Map showing approximate ranges of *Chrysemys* turtles. A red star marks McLeod County MN, the collection site of turtle R12L10, which was used for the assembly presented here. Adapted from: https://en.wikipedia.org/wiki/Wikipedia:Featured_picture_candidates/Painted_turtle_native_range#Wikipedia:Featured_picture_candidates/Painted_turtle_native_range

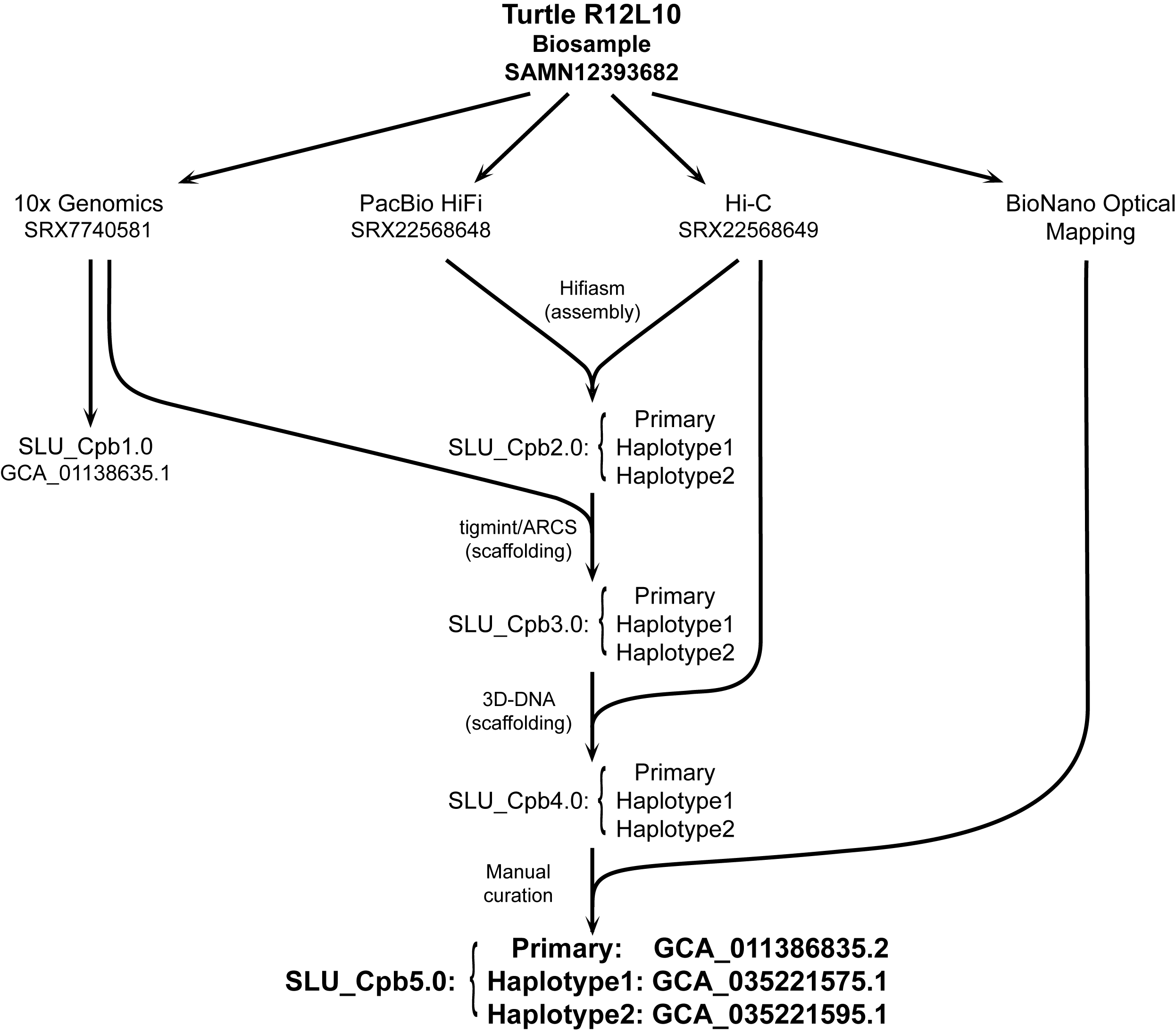

**Legend to Additional Figure S2: Flowchart describing procedures of genome assembly in this study.** An initial *C. picta bellii* genome assembly was generated using 10x Genomics linked reads (ASM1138683v1). Additional PacBio HiFi long reads, Hi-C conformation capture sequence data, and Bionano optical genome maps were generated from the same individual turtle. Different *de novo* genome assembly tools were used for de novo genome assembly using PacBio HiFi reads. The assembly generated by Hifiasm with the integration of Hi-C reads yielded the best metrics in assemblies, which were used for subsequent scaffoldings. The first round of scaffolding for the preliminary assemblies was performed with the 10x Genomics linked reads using Tigmint. The second round of scaffolding used Hi-C sequencing reads by 3D-DNA. The primary assembly was aligned with BioNano optical maps, and major discrepancies were corrected manually.

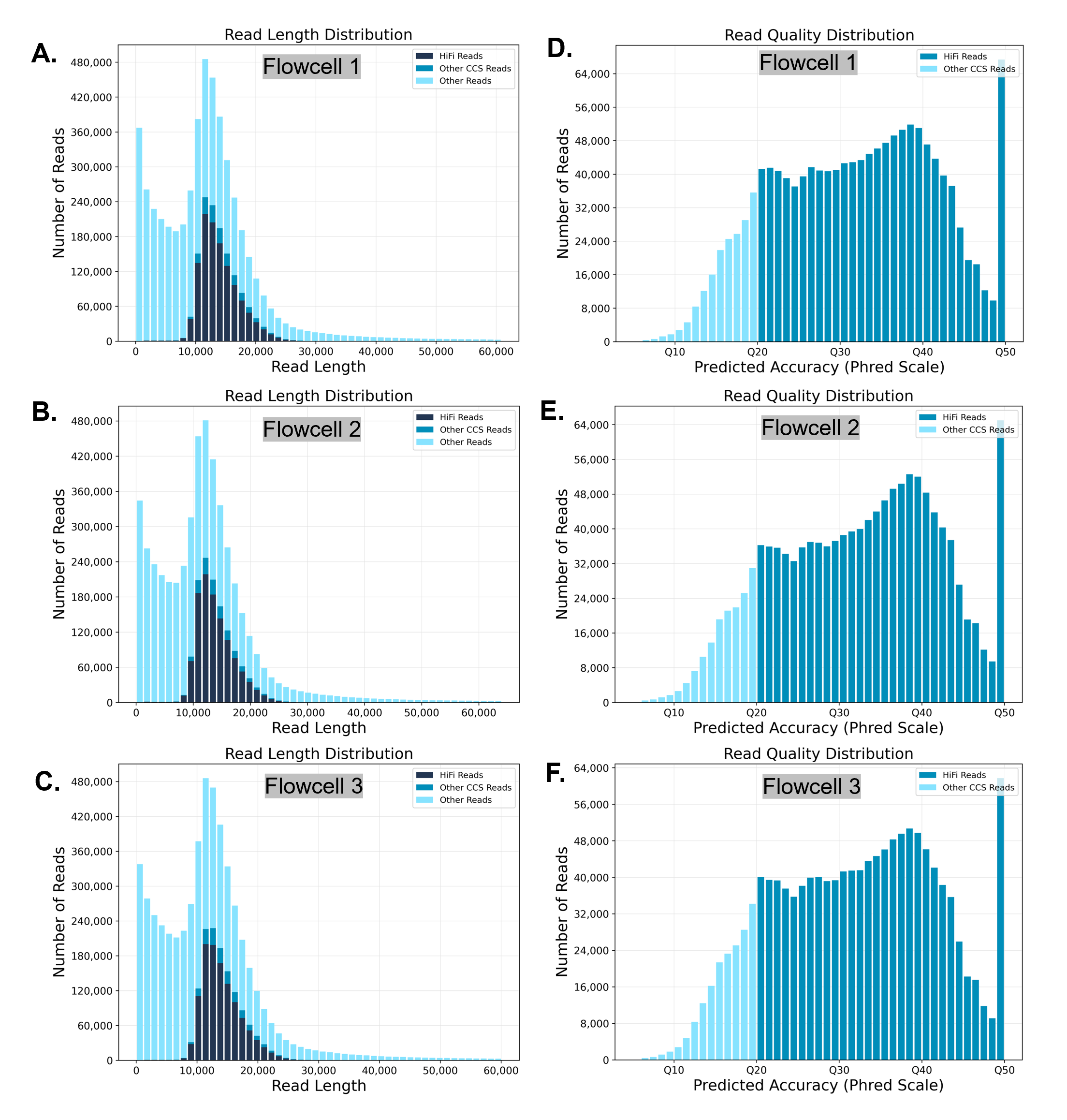

**Legend to Additional Figure S3: Read length and quality distribution of PacBio HiFi sequencing data. (A-C):** Histograms displaying the distribution of read lengths for PacBio HiFi sequencing data from the turtle DNA sample of three flowcells, respectively. HiFi reads (high-accuracy circular consensus sequences); Other CSS reads (circular consensus sequences that do not meet HiFi quality thresholds); and Other reads (non-consensus or lower-quality reads). (**D-F):** Histogram showing the distribution of read quality (Phred score) for PacBio HiFi sequencing data and other CSS reads from the turtle DNA sample of three flowcells.

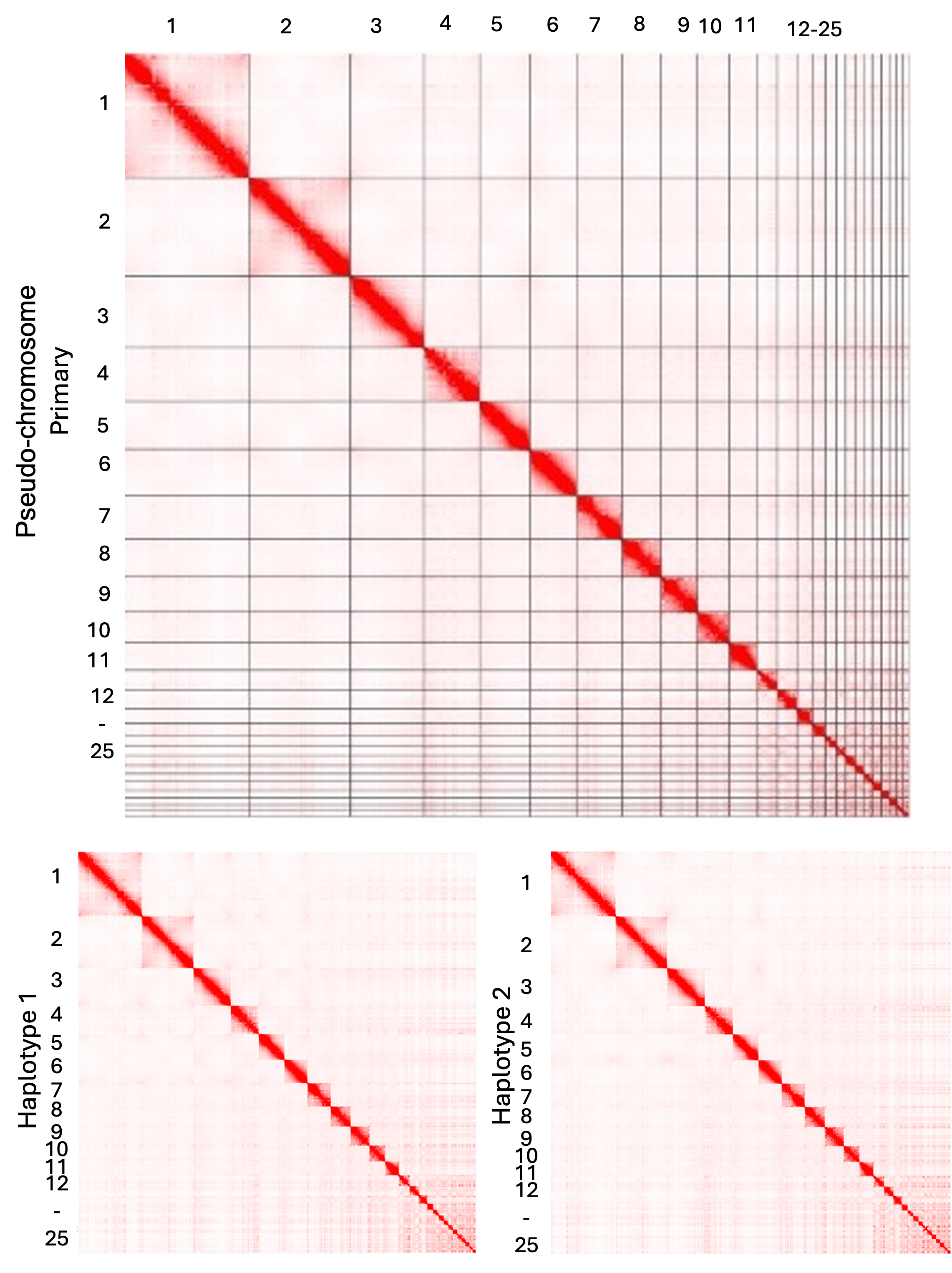

**Legend Additional Figure S4: Hi-C contact map visualizing chromosomal interactions.** A red point depicts each Illumina read and its corresponding mate read; clusters of points denote higher contact frequency, suggesting closer spatial proximity in the genome. The maps display interaction patterns for three assemblies: **(A)** Primary Assembly. **(B)** Haplotype 1. **(C)** Haplotype 2. The contact maps were generated using Juicebox v1.11.08.

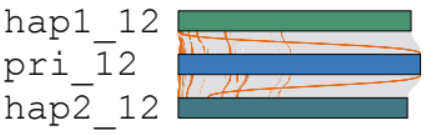

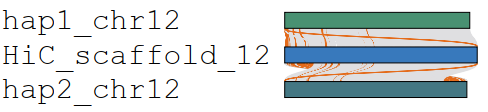

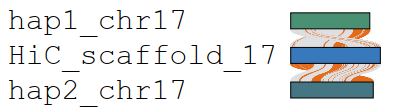

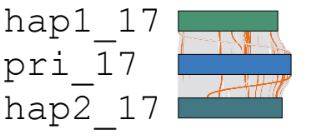

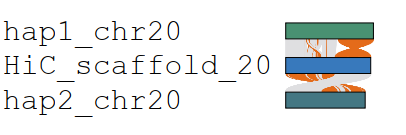

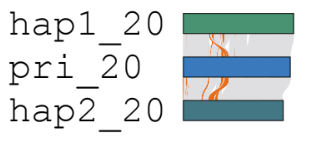

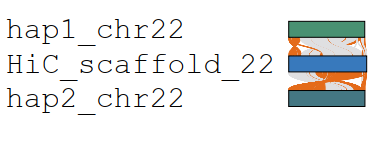

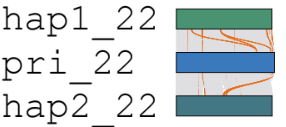

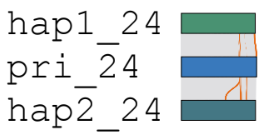

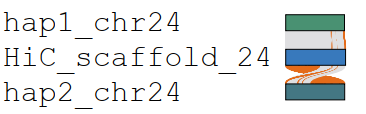

Hap1- Chr. 12

Hap2 - Chr. 12

Primary - Chr. 12

Before

After

Hap1- Chr. 12

Hap2 - Chr. 12

Primary - Chr. 12

Hap1- Chr. 17

Hap2 - Chr. 17

Primary - Chr. 17

Hap1- Chr. 17

Hap2 - Chr. 17

Primary - Chr. 17

Hap1- Chr. 20

Hap2 - Chr. 20

Primary - Chr. 20

Hap1- Chr. 20

Hap2 - Chr. 20

Primary - Chr. 20

Hap1- Chr. 24

Hap2 - Chr. 24

Primary - Chr. 24

Hap1- Chr. 24

Hap2 - Chr. 24

Primary - Chr. 24

Hap1- Chr. 22

Hap2 - Chr. 22

Primary - Chr. 22

Hap1- Chr. 22

Hap2 - Chr. 22

Primary - Chr. 22

**Additional Figure S5: Examples of manual corrections in chromosome assemblies using Hi-C contact maps and BioNano optical mapping.** The sequences of three assemblies were aligned to identify major discrepancies among them using NGenomeSyn. In the "Before" panel (left), Hi-C contact maps and BioNano optical maps highlight issues such as incorrect orientations and inversions. These errors were manually corrected, and the revised assemblies were subsequently aligned to illustrate the improvements in the "After" panel (right). Graphical alignments between haplotypes and scaffolds for five chromosomes are shown. Chromosomes/scaffolds are shown as bars; discontinuous regions of homology between them are shown in brown. After manual curation, regions of discordance are reduced (right).

**A.**

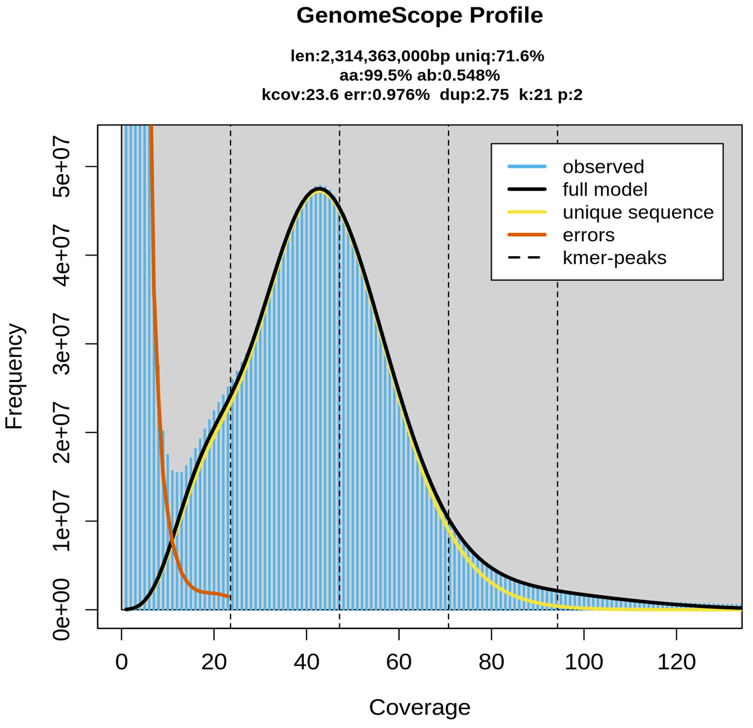

**B.**

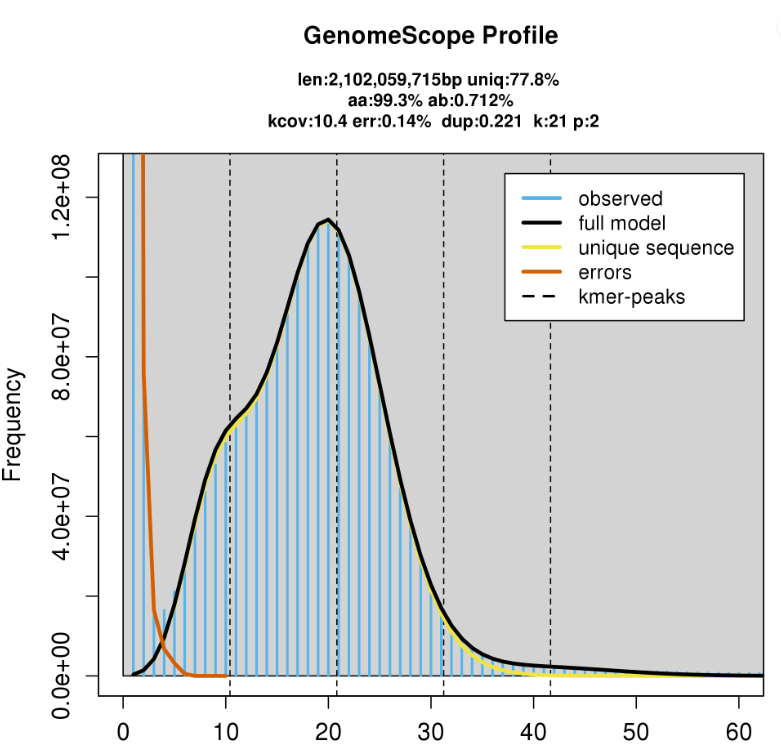

**Additional Figure S6: The K-mer distribution of different sequencing reads of turtle genomic DNA. (A):** The K-mer distribution of 10X Genomics Illumina paired-end reads. (**B):** The K-mer distribution of PacBio HiFi reads. The K-mer distribution in each sample was calculated by GenomeScope2.0 (Ranallo-Benavidez et al. 2020) based on k value of 21. Blue bars represent the observed k-mer distribution. The black line represents the predicted distribution of k-mer frequencies based on genome properties. The presence of a single peak indicates that the diploid genome has a low heterozygosity. A good fit in both graphs supports high reliability of the two types of sequencing data generated by this study.

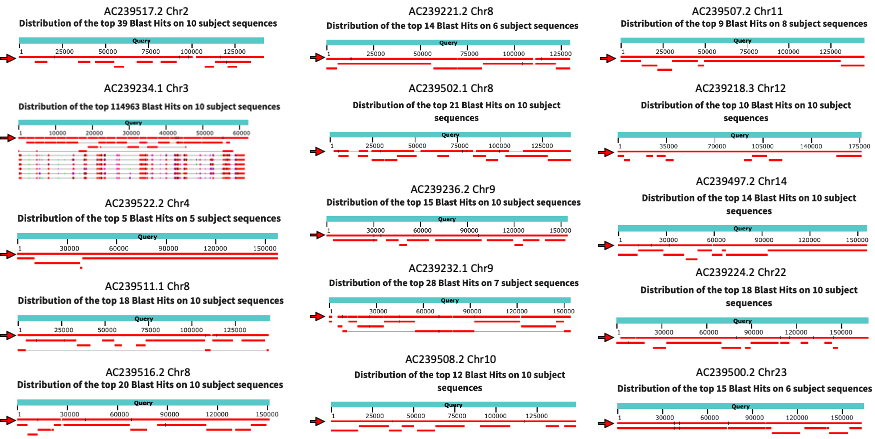

**Additional Figure S7: Graphic displays of BLAST localization of representative BACs to *C.picta bellii* genome assemblies.** Select examples show greater contiguity in our chromosome-level assembly. In each panel, a BAC accession number and chromosomal location are given. Red bars that are marked with an arrow represent alignments with the *C. picta bellii* assembly from this study, whereas the others represent RCT428 (CPI3.0.4) contigs.

**A:**

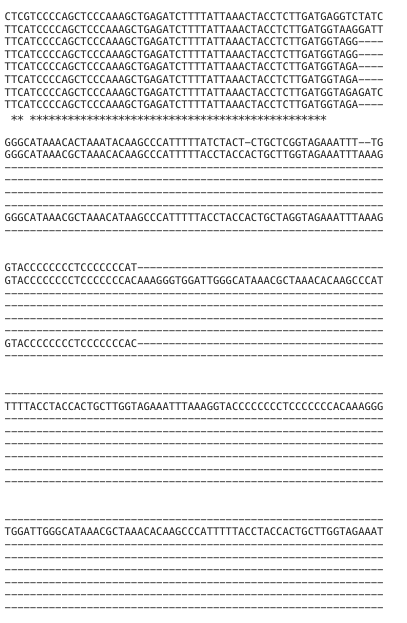

**tRNAPro (rev. complement)**

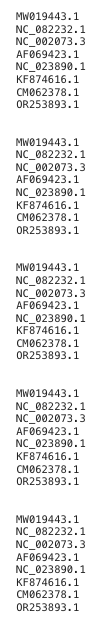

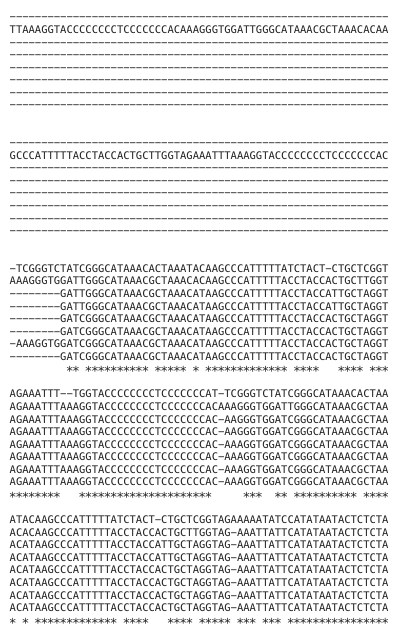

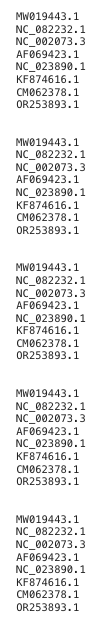

**VNTR1A: 59bp	VNTR1B: 33bp**

**B**

**Emydinae**

**Deirochelyinae**

**Emydinae**

**Deirochelyinae**

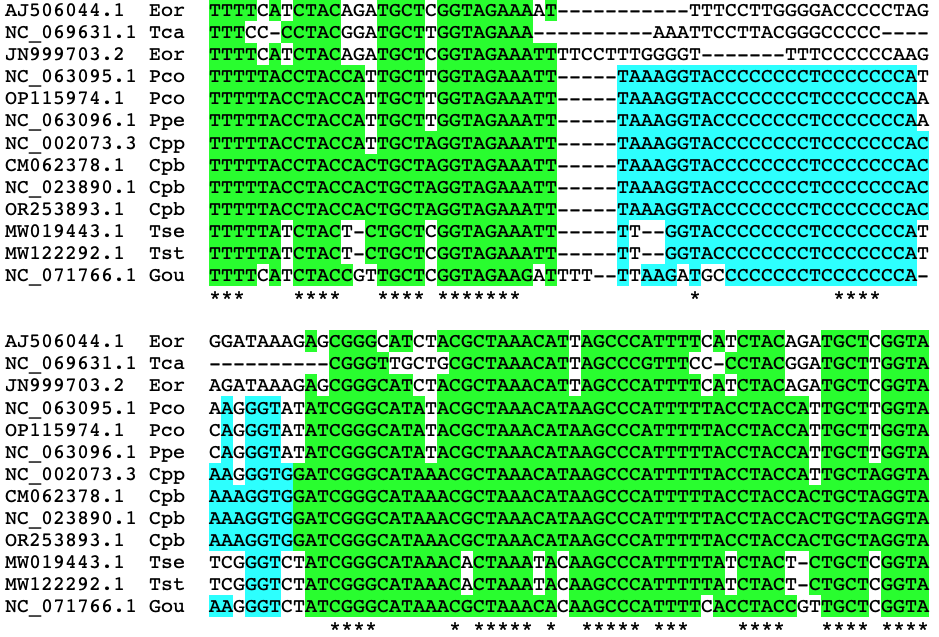

**Legend to Additional Figure S8: Mitochondrial VNTR1 repeats.** A. Chrysemys-Trachemys mitochondrial VNTR repeats. Different *Chrysemys* turtles possess differing numbers of VNTRA and VNTRB repeats. Repeat number is variable even among *C. picta bellii* mitochondria. Repeat lengths are as follows: ABABABABABA: Chrysemys Dorsalis (NC_082232); ABABA: Trachemys s. elegans MW019443; *Chrysemys p. bellii* CM062378; ABA: *Chrysemys picta* NC_002073, AF069423; *Chrysemys p. bellii* NC_023890, KF874616; OR253893. B. VNTR1A, VNTR1B conservation in Emydidae. Alignments of a VNTR1B flanked by the 3’ end of one VNTR1A, and the 5’ end of a second VNTR1A. The alignments of sequences derived from members of both subfamilies of the turtle family Emydidae demonstrate that VNTR1A is more conserved than is VNTR1B. Species abbreviations: Emydinae: Eor: Emys orbicularis; Tca: *Terrapene carolina*. Deirochelyinae: Pco: *Pseudemys concinna*; Ppe: *Pseudemys peninsularis*; Cpp: *Chrysemys picta picta*; Cpb: Chrysemys picta bellii; Gou: *Graptemys ouachitensis*; Tst: *Trachemys scripta troostii*; Tse: *Trachemys scripta elegans*.

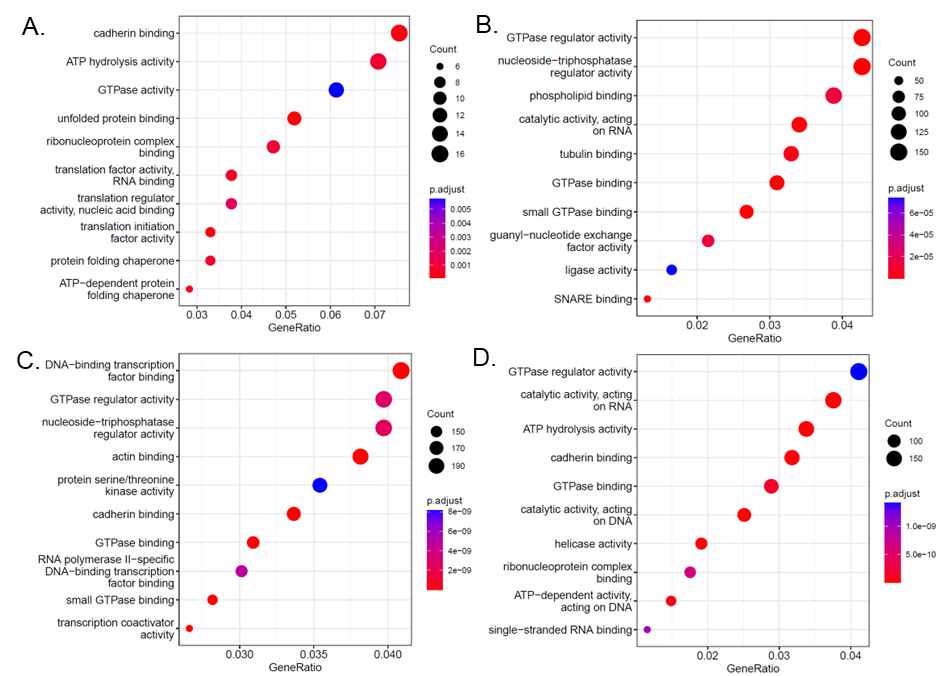

**Additional Figure S9: Gene Ontology (GO) analyses reveal enriched biological functions of shared and tissue specific transcript isoforms. (A).** Shared transcript isoforms. (B) Transcript isoforms specific to hatchling brain. **(C)**. Transcript isoforms specific to hatchling carapace. **(D)** Transcript isoforms specific to testis.

**Additional Table Legends:**

**Legend to Additional Table S1: NCBI Accession numbers for sequencing data and assemblies generated in this study.** Biosample, Bioproject, Sequence Run Accession, Assembly, Accession, and URLs are listed for the datasets that were used in generating the SLU_Cpb5.0 genome assembly.

**Legend to Additional Table S2: Summary statistics for three PacBio HiFi flowcells**

**Legend to Additional Table S3: Genome assembly statistics for intermediate assemblies**

**Legend to Additional Table S4: Estimation of genome size and heterozygosity using GenomeScope:** GenomeScope statistics for Illumina (10x Genomics) and PacBio HiFi sequence datasets.

**Additional Table S5: RNAseq datasets used in this study. The table gives alignment rates to the previous (BioNano-3.0.4) and current (SluCpb5.0) genomes.** Discordant pairs have unique alignments but do not match paired-end expectations.

**Legend to Additional Table S6: Localization of BAC sequences on SLU_Cpb2.0 genome assembly.** The Table lists 51 bacterial artificial chromosomes (BACs) that were localized to C.picta bellii metaphase chromosomes by fluorescent in situ hybridization (FISH). All but CHY3-16H12 (Accession #AC239505.2) are analyzed in [1]; BACs used in [2] have not been given accession numbers, and therefore could not be analyzed. FISH chromosome localizations on the CPI3.0.4 genome and on the SLU_Cpb2.0 genome assembly are listed. Differences in chromosome assignments that are attributable to similarly sized metaphase chromosomes are shown in blue; differences in chromosome orientation are shown in yellow. Discrepancies in chromosome assignment between FISH and sequencing studies are shown in red. BACs shown in purple are difficult to localize on the SLU_Cpb2.0 assembly due to multiple copies of sequences, or to unordered BAC sequences. BACs whose alignments are shown in **Additional Figure 7** are shaded.

**Legend to Additional Table S7: *Chrysemys* mitochondrial sequence identities, divergence**

Percent identities, shown above the diagonal, highlighted in GREEN, were obtained from NCBI pairwise BLAST alignments of Chrysemys mitochondrial genomes. Percent divergences, shown below the diagonal and highlighted in BLUE, were calculated by subtracting percent identities from 100%. Korean *Chrysemys picta bellii* population [3] came from the Midwest, not the Northwest.

**Legend to Additional Table S8: *Chrysemys* mitochondrial polymorphisms**. The table shows polymorphisms relative to the R12L10 mitochondrial genome (NCBI Accession # CM062378. Column 1 gives the nucleotide position of the polymorphism; Column 2 shows the polymorphism; del=deletion. Column 3 shows the turtle mitochondrial genome in which the polymorphism occurred: Abbreviations: Cd: *C. dorsalis* NC_082232.1; Cpp: *C.picta picta* NC_002073.3; Cpb(WA): *C.picta bellii* from Washington state, NC_028390.1; Cpb(K): *C.picta bellii* from South Korea, OR253893.1. Column 4 lists the mitochondrial gene in which the polymorphism occurred: Abbreviations: transmemb. = transmembrane domain; comp = complementary strand. Outside and inside refer to transmembrane orientation. Column 5 lists amino acid changes that result from the polymorphisms. Columns 6 & 7 depict whether the amino acid changes are conservative (predicted to be less likely to affect protein function) or radical (more likely to alter protein function). Conservation was evaluated using BLOSUM62 scores, which are based on observed amino acid substitution frequencies in aligned conserved sequences [4], or from a table of predicted effects of amino acid substitutions on protein folding in silico [5]. Aledo and Aledo used two criteria: Effects on folding energy and amino acid polarity or volume. Conservative changes are shaded in BLUE; amino acid changes that have radical effects on folding energy but not on polarity-volume are shaded in GREEN; those with radical effects on polarity-volume but not on folding energy are shaded in PINK, and those with radical effects on both criteria are shaded in RED. Several amino acid substitutions are highlighted as non-conservative by both BLOSUM62 and in silico modeling; they may underlie physiological differences between *Chrysemys* populations. Amino acid changes that are within three amino acids from a transmembrane domain junction, as predicted by DeepTMHMM (https://dtu.biolib.com/DeepTMHMM) are listed in bold typeface. Mitochondrial sequences that were amplified for phylogenetic predictions by [6] and [7] are shaded in gray.

**Legend to Additional Table S9: Statistics of PacBio Isoseq sequencing for 12 tissues.** Numbers of reads, and total sequence obtained is given for each of the tissues that were used for PacBio isoseq long-read RNAseq.

**Legend to Additional Table S10: Types of transcript isoforms identified in each tissues based onPacBio Iso-seq data.**

**Legend to Additional Table S11: Presence of absence of transcript isoforms in each tissues based on PacBio Iso-seq data.**

**Legend to Additional Table S12: Summary of RNAseq differential expression data with the new assembly.**

**Legend to Additional Table S13: Comparison of DESeq2 results for RNA-seq data in telencephalon between genome assembly SLU_Cpb5.0 and CPI3.0.4.**

**Additional Table S14: Comparison of DESeq2 results for RNA-seq data in ventricle between genome assembly SLU_Cpb5.0 and CPI3.0.4**

**Additional References:**

1.	Badenhorst D, Hillier LW, Literman R, Montiel EE, Radhakrishnan S, Shen Y, Minx P, Janes DE, Warren WC, Edwards SV, Valenzuela N: **Physical Mapping and Refinement of the Painted Turtle Genome (Chrysemys picta) Inform Amniote Genome Evolution and Challenge Turtle-Bird Chromosomal Conservation.** *Genome Biol Evol* 2015, **7:**2038-2050.

2.	Lee LS, Navarro-Dominguez BM, Wu Z, Montiel EE, Badenhorst D, Bista B, Gessler TB, Valenzuela N: **Karyotypic Evolution of Sauropsid Vertebrates Illuminated by Optical and Physical Mapping of the Painted Turtle and Slider Turtle Genomes.** *Genes (Basel)* 2020, **11**.

3.	Park J, Park SM, Choi JH, Sung HC, Lee DH: **Complete mitochondrial genome of the western painted turtle (*Chrysemys picta bellii,* Testudines: Emydidae) in Korea.** *Mitochondrial DNA B Resour* 2023, **8:**1316-1319.

4.	Henikoff S, Henikoff JG: **Amino acid substitution matrices from protein blocks.** *Proc Natl Acad Sci U S A* 1992, **89:**10915-10919.

5.	Aledo P, Aledo JC: **Proteome-Wide Structural Computations Provide Insights into Empirical Amino Acid Substitution Matrices.** *Int J Mol Sci* 2023, **24**.

6.	Starkey DE, Shaffer HB, Burke RL, Forstner MR, Iverson JB, Janzen FJ, Rhodin AG, Ultsch GR: **Molecular systematics, phylogeography, and the effects of Pleistocene glaciation in the painted turtle (Chrysemys picta) complex.** *Evolution* 2003, **57:**119-128.

7.	Jensen EL, Govindarajulu P, Russello MA: **When the shoe doesn’t fit: applying conservation unit concepts to western painted turtles at their northern periphery.** *Conservation Genetics* 2014, **15:**261-274.
